# Colonization by orchid mycorrhizal fungi primes induced systemic resistance against a necrotrophic pathogen

**DOI:** 10.1101/2024.02.07.579401

**Authors:** Galih Chersy Pujasatria, Chihiro Miura, Katsushi Yamaguchi, Shuji Shigenobu, Hironori Kaminaka

**Affiliations:** The United Graduate School of Agricultural Sciences, Tottori University, Tottori, Japan; Faculty of Agriculture, Tottori University, Tottori, Japan; Functional Genomics Facility, National Institute for Basic Biology Core Research Facilities, Okazaki, Japan; Unused Bioresource Utilization Center, Tottori University, Tottori, Japan

**Keywords:** *Bletilla striata*, defense priming, *Dickeya fangzhongdai*, induced systemic resistance (ISR), mycorrhizal fungi, necrotrophic pathogen, orchids, *Serendipita vermifera*, *Tulasnella calospora*

## Abstract

Orchids and arbuscular mycorrhiza (AM) plants evolved independently and have different structures and fungal partners, but they both facilitate nutrient uptake. Orchid mycorrhiza (OM) supports orchid seed germination, but unlike AM, its role in disease resistance of mature plants is largely unknown. Here, we examined whether OM induces systemic disease resistance against a necrotrophic pathogen in a similar fashion to AM. We investigated the priming effect of mycorrhizal fungi inoculation on resistance of a terrestrial orchid, *Bletilla striata*, to soft rot caused by *Dickeya fangzhongdai*. We found that root colonization by a compatible OM fungus primed *B. striata* seedlings and induced systemic resistance against the infection. Transcriptome analysis showed that priming was mediated by the downregulation of jasmonate and ethylene pathways and that these pathways are upregulated once infection occurs. Comparison with the reported transcriptome of AM fungus–colonized rice leaves revealed similar mechanisms in *B. striata* and in rice. These findings highlight a novel aspect of commonality between OM and AM plants in terms of induced systemic resistance.

**Highlight:** Colonization by a compatible mycorrhizal fungus primes induced systemic resistance against a necrotrophic pathogen in a terrestrial orchid, *Bletilla striata*, by regulating jasmonate and ethylene pathways, similar to arbuscular mycorrhizal plants.

## Introduction

Mycorrhiza is one of the oldest symbioses, in which plant roots are colonized by fungi forming a specialized structure inside or outside root cells where nutrient uptake and transfer occur (Smith and Read, 2008). The most common type of mycorrhiza, arbuscular mycorrhiza (AM), is found in almost 80% of flowering plants (Brundrett and Tedersoo, 2018) and appeared in the rooting system at the latest during the Carboniferous (Strullu-Derrien *et al*., 2018). During mycorrhizal colonization, AM fungi transfer nitrogen, phosphate and water to the host plants (Parniske, 2008; Kakouridis *et al*., 2022). In return, plants provide carbon compounds to the fungi. The extent of each of these physiological functions often differs among mycorrhizal types and plant species. On the basis of these differences, several other mycorrhizal types are distinguished, including ectomycorrhiza, orchid mycorrhiza (OM), and ericoid mycorrhiza. OM is unique in that the fungal structures formed inside orchid cells are transient and are gradually degraded; this phenomenon is most prominent during early seed germination (Selosse *et al*., 2017). Mycorrhizal associations in orchids are obligatory during early germination, although in green, leafy species their extent decreases with time due to the plants’ autotrophic nature (Cameron *et al*., 2008; Girlanda *et al*., 2011).

Orchidaceae have evolved endomycorrhizas with Basidiomycota or occasionally Ascomycota (Fracchia *et al*., 2016; Strullu-Derrien *et al*., 2018). The dominant group, *Rhizoctonia-*like fungi, includes three main genera—*Ceratobasidium*, *Sebacina/Serendipita*, and *Tulasnella* (Smith and Read, 2008)—while some lower taxa of subfamily Epidendroideae are sometimes associated with ectomycorrhizal fungi (McKendrick *et al*., 2000; Bidartondo *et al*., 2004; Zahn *et al*., 2023) or even wood-rotting, mushroom-forming fungi (Umata *et al*., 2013). Both OM and AM are involved in nitrogen, carbon, and phosphate transfer (Cameron *et al*., 2007, 2008; Kuga *et al*., 2014; Fochi *et al*., 2016) and have common integral molecular mechanisms (Miura *et al*., 2018). It was recently discovered that gibberellin inhibits both OM and AM colonization, but it inhibits seed germination only in orchids (Miura *et al*., 2024). These findings further encourage the exploration of the commonalities between OM and AM, including ecophysiological aspects such as disease resistance.

Two distinct defense responses occur upon pathogen infection: an early, local response at the infection site and a systemic response at distal sites (David *et al*., 2019); the latter includes systemic acquired resistance (SAR) (Ross, 1966) and induced systemic resistance (ISR) (van Peer *et al.,* 1991; Wei *et al.,* 1991). Systemic acquired resistance occurs when pathogens directly infect leaves and involves mainly salicylic acid, whereas ISR is induced by beneficial microbes (bacteria, endophytic fungi, and mycorrhiza) interacting with roots and involves jasmonate and ethylene (Vlot *et al*., 2021). During ISR, the plant enters an alert state called priming, allowing enhanced resistance once infection occurs, mainly against necrotrophic pathogens (Pieterse *et al*., 2014). Primed plants often show no visible changes prior to infection; thus, ISR is best studied in plants challenged with pathogens, which allows changes at the cellular level between non-primed and primed plants to be compared. Due to the diversity of microbes interacting with roots, the molecular mechanisms of ISR are more diverse than those of SAR because different microbe species induce the accumulation of different signaling compounds (Haney *et al*., 2018; Dreischhoff *et al*., 2020). ISR induced by an AM fungus in *Medicago truncatula* increases the defense response against *Xanthomonas campestris* (Liu *et al*., 2007), whereas *Solanum lycopersicum* has a similar response against *Fusarium oxysporum* (Wang *et al*., 2022a).

Almost all types pathogens attack orchids, including bacteria (Keith *et al*., 2005; Suharjo *et al*., 2014; Khamtham and Akarapisan, 2019), fungi (Silva and Pereira, 2007; Lopes *et al*., 2009; Srivastava *et al*., 2018; Suwannarach *et al*., 2018), and viruses (Fogell *et al*., 2019; Tsai *et al*., 2022). Very little is known about ISR in orchids; to the best of our knowledge, only two studies have reported the occurrence of ISR and its role in alleviating soft rot (Wu *et al*., 2011; Ye *et al*., 2019). Thus, enhancing our knowledge on ISR in orchids will give clues on the consequences of evolving mycorrhizal association.

On the basis of the arguments that colonization by AM fungi primes host defense responses against pathogens (Marquez *et al*., 2018) and major molecular components of AM symbiosis signaling are also present in OM (Miura *et al*., 2018), here we hypothesized that colonization by OM fungi (OMF) also causes ISR in orchids just as in AM plants. We used a necrotrophic, pectinolytic Gram-negative bacterium known to cause leaf soft rot in orchids (Cating and Palmateer, 2011; Joko *et al*., 2014; Wei *et al*., 2021). As recommended by Eck *et al*. (2022), we also investigated the relationship between mycorrhizal colonization rate and ISR. We focused on a generalist orchid species, *Bletilla striata* (Orchidaceae, tribe Arethuseae), because it is relatively easy to grow and it associates with OMF from different genera (Yamamoto *et al*., 2017; Miura *et al*., 2019). Using two OMF species previously reported to associate with *B. striata* (Miura *et al*., 2019; Fuji *et al*., 2020), we evaluated ISR and its molecular regulation in leaves through transcriptomic analysis.

## Materials and Methods

### Plant and fungal materials

*Bletilla striata* Rchb.f. ‘Murasakishikibu’ plants were purchased from a local nursery in Japan (Yamamoto *et al*., 2017). For seed production, flowers were self-pollinated and allowed to set seeds for 6 months (until dehiscence). Seeds were randomly picked from capsules harvested between 2017 and 2020. In a pathogen inoculation assay, aside from *B. striata*, *Arabidopsis thaliana* (L.) Heynh. ecotype Columbia (Col-0) and *Nicotiana benthamiana* were also used.

Our earlier studies showed that several strains of *Tulasnella* and *Serendipita vermifera* form OM associations with *B. striata* during seed germination (Yamamoto *et al*., 2017; Miura *et al*., 2019; Fuji *et al*., 2020). Among those fungi, we chose *T. calospora* (Boud.) Juel MAFF305805 (coded as T.cal05) and *S. vermifera* (Oberw.) P. Roberts MAFF305830 (coded as S.ver30).

### Symbiotic seed germination

In symbiotic seed germination, *S. vermifera* and *T. calospora* were cultured in agar (1.5 g/L) containing 2.5 g/L oatmeal (Becton Dickinson, Sparks, MD, USA) until maximum hyphal growth. *Bletilla striata* seeds were surface-sterilized in 3 mL of 5% NaOCl containing 5 µL Tween-80 for 2 min and were sown directly on top of the mycelia. The plates were incubated at 25°C for 2 weeks.

### Asymbiotic seed germination and direct root inoculation of OMF

In asymbiotic seed germination, *B. striata* seeds were surface-sterilized as above and sown on solid half-strength Murashige–Skoog (MS) medium supplemented with 20 g/L sucrose, 20 g/L banana homogenate, 1 g/L tryptone, 3 g/L activated charcoal, MS vitamin mixture (pH 5.8), and 0.8 g/mL agar. Seedlings were grown for 8 weeks until formation of true leaves and roots. For the first subculture, the plantlets were transferred into half-strength P668 medium (Phytotech Labs, Lenexa, KS, USA) supplemented with 20 g/L sucrose, 0.1 g/L tryptone, 0.2 mg/L naphthalene acetic acid, 0.5 mg/L 6-benzylaminopurine, MS vitamin mixture (pH 5.8), and 0.8 g/mL agar for another 6–8 weeks. Seedlings with at least two true leaves and new roots longer than 2 cm were transferred into small pots filled with a mixture of *akadama* soil, *kanuma* soil, and vermiculite (4:4:1 volume ratio) under a 16 h light/8 h dark photoperiod at 25°C for another 2 weeks for hardening (Fig. S1). *Serendipita vermifera* and *T. calospora* were cultured in YEPG liquid medium containing 3 g/L yeast extract, 3 g/L peptone, and 20 g/L glucose for 4 weeks until sufficient mycelial growth. The mycelial mass was harvested and homogenized in distilled water (4 mL/g mycelial mass) using a homogenizer (Nissei, Osaka, Japan). A 2-mL aliquot of the mycelial suspension was inoculated beneath each seedling.

### Visualization and quantification of OMF colonization in protocorms and roots

Root colonization is defined as the presence of pelotons, both intact and degraded (Yamamoto *et al.,* 2017). Protocorms and roots were fixed in 70% ethanol and cleared in 5% KOH at 90°C for 1 h (Peterson *et al*., 2004). To evaluate colonization, the protocorms were stained with 5% black ink (Sheaffer, Fort Madison, IA, USA) in 5% acetic acid for 10 min (Vierheilig *et al*., 1998). Protocorms were photographed under a light microscope (BX53; Olympus, Tokyo, Japan) equipped with a digital camera (DP27; Olympus). By using ImageJ v.1.53a, pelotons present in protocorms were counted with the multipoint tool and root colonization was measured with the segmented line tool. Root colonization was quantified as described for AM plants (McGonigle *et al*., 1990) with modifications: each piece of root was divided into imaginary 1-mm segments. Each segment was further divided horizontally into four subsegments, especially the middle part by the vascular bundle. The ratio of cortical colonization within each segment was scored as 0 (no OMF colonization on either side), 0.25 (one-quarter of the segment is colonized), 0.5 (only one side is fully colonized), 0.75 (three-quarters of the segment is colonized), or 1 (both sides are colonized). The colonization rate (CR) for a single root was calculated through the colonization index (CI) using the following equations:

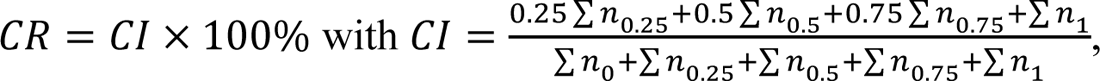

where *i* is a positive integer from 0 to 4, *n_i/_*_4_ is the number of segments colonized at a particular level, and *N* is the total number of segments in one root. This index can vary from 0 to 1. The final CR of a seedling was then expressed as the arithmetic average value of CR obtained from all roots. The detailed method is available at Zenodo (https://doi.org/10.5281/zenodo.10464582).

### Pathogen inoculation

*Erwinia chrysanthemi* MAFF311045, which was isolated from *Phalaenopsis* (Suharjo *et al*., 2014), was cultured in Luria–Bertani (LB) medium overnight. Bacterial suspension at several concentrations and OD_600_ = 0.0025 was mixed with 0.01% Tween-20 (10:1 volume ratio). Two leaves per seedling of *B. striata*, *A. thaliana*, and *N. benthamiana* were used. For each leaf, one small drop (3 µL) was placed on the syringe-wounded adaxial surface. During infection, high humidity was maintained by flooding the planting tray with water and covering the plantlets with a moist plastic lid. Symptoms were observed and leaf samples were collected 3 days after inoculation. Infected leaves (diameter 4 mm) were excised at the site of initial infection and macerated in 1 mL liquid selective enhancement medium containing 3.75 g/L MgSO_4_·7H_2_O, 1 g/L (NH_4_)_2_SO_4_, 1 g/L K_2_HPO_4_, 0.2 mL/L 5N NaOH, and 1.7 g/L pectin at pH 7.2 (Wako Pure Chemicals, Osaka, Japan) for 24 h at 28°C (Hélias *et al*., 2012). Because of the ability of this pathogen to produce indigoidine (Lee and Yu, 2006; Alič *et al*., 2019; Wei *et al*., 2021), we used a selective nutrient agar medium containing crystal violet, 70 mg/mL glutamine, and 4 mg/mL MnCl_2_·2H_2_O at pH 6.5–6.8. Plates were incubated at 28°C for 1 day and bluish colonies were selected for counting.

### Leaf symptoms, indicators of photosynthetic damage, and peroxide content

To visualize the necrotic area, leaves were immediately placed into 0.05% trypan blue, stained at 37°C overnight, and destained in absolute ethanol for another night. To visualize peroxide distribution around the infected area, infected leaf samples were directly incubated in 1 mg/mL diaminobenzidine at 37°C overnight in the dark. The leaves were fully decolorized in several changes of 70% ethanol at 70°C for 1 h and stored in 30% glycerol until observation.

Photosynthetic quantum yield was measured with a miniPPM-300 photosynthesis meter (EARS, Wageningen, Netherlands) after dark preconditioning for at least 30 min. Subsequently, individual leaves were weighed, and chlorophyll was extracted with 1 mL dimethyl sulfoxide per leaf overnight. Quantification of total chlorophyll content was based on extract absorbance at 649 and 665 nm and an equation suggested by Wellburn (1994). The values expressed in µg/mL solvent were converted into µg/g leaf fresh weight.

The total peroxide content of leaves was measured by luminol chemiluminescence catalyzed by Co^2+^ (Pérez and Rubio, 2006). Samples were frozen in liquid nitrogen, ground in 0.5 mL 5% trichloroacetic acid, and centrifuged at 13,000 rpm and 4°C for 10 min. The supernatant was supplemented with 5% polyvinylpyrrolidone and diluted 100-fold with water. An aliquot of the diluted extract (20 µL) was added to 1 mL of diluted luminol– cobalt mixture (1 g/L luminol and 0.7 g/L CoCl_2_·6H_2_O in carbonate buffer pH 10.2, incubated for at least 1 h, and diluted 10-fold before use). Chemiluminescence was measured 5 s after the addition of luminol in a Gene Light GL-220 luminometer (Microtec Nition, Chiba, Japan). The values were expressed as log_10_ (relative light unit)/g fresh weight.

### RNA extraction and RNA-sequencing

Aside from control, only *T. calospora-*colonized seedlings, regardless of their infection status, were used for RNA extraction and sequencing. Each replicate contained materials collected from at least five individuals. Total leaf RNA was extracted using a Total RNA Extraction Kit (Plant) (RBC Bioscience, Taipei, Taiwan) according to the manufacturer’s protocol. Residual genomic DNA was removed with RNase-free DNase I (Toyobo, Osaka, Japan). The quality and quantity of the total RNA were checked using a Qubit RNA HS Assay Kit and Qubit 2.0 Fluorometer (Thermo Fisher Scientific, Waltham, MA, USA).

RNA-seq library construction from the total RNA using an MGIEasy RNA Directional Library Prep Set (MGI, Shenzhen, China) and sequencing with strand-specific and paired-end reads (150 bp) using the DNBSEQ-T7RS sequencing platform were performed by Genome-Lead Co. (Takamatsu, Kagawa, Japan).

### Transcriptome analysis

The obtained raw reads were filtered using Fastp v.0.23.2 (Chen *et al*., 2018) to remove low-quality reads (<Q30) and adapter sequences. Reads were mapped to *de novo* assembled transcript sequences (Miura *et al*., 2018) using Bowtie2 v.2.5.2 (Langmead and Salzberg, 2012). Differentially expressed genes (DEGs) were analyzed using the *edgeR* package. Library size was corrected using the trimmed mean of M value method and fitted using the *glmTreat* (filtered at logFC 1.5) (Smyth *et al*., 2018). Gene Ontology (GO) enrichment was analyzed in the *topGO* package (Alexa and Rahnenfuhrer, 2023) using the *elim* parameter and Fisher exact test, and the *p*-values were adjusted with the Benjamini– Hochberg method. The false discovery rate (FDR) threshold was set at 0.05. Genes of *B. striata* orthologous to rice genes were identified using SonicParanoid (Cosentino and Iwasaki, 2019) with default parameters. The rice genome assembly (Os-Nipponbare-Reference-IRGSP-1.0; Kawahara *et al.,* 2013) were retrieved from Ensembl Plants (https://plants.ensembl.org/index.html). Additional genes involved in jasmonate and ethylene biosynthesis and phosphate metabolism were retrieved from rice transcriptomic data (Campo and San Segundo, 2020) and used as queries for local TBLASTX against *B. striata* genome assembly. Only *B. striata* genes with an E-value less than 10^−5^ were selected. The whole TBLASTX process was conducted in Genetyx v.15 software (Genetyx, Tokyo, Japan). For visualization, a heatmap was generated in the RStudio v.4.1.2 package *Complexheatmap* (Gu *et al*., 2016), and an UpSet plot was generated manually using CorelDRAW 2021 v.23.1.0.389 (Alludo, Ottawa, ON, Canada).

### Statistical analysis

For protocorm symbiotic cell count, root colonization rate, *D. fangzhongdai* titer, photosynthetic quantum yield, chlorophyll content, and peroxide content data, whenever the data did not meet the normality assumption, logarithmic data transformation was used to enable analyses with parametric tests according to the corresponding data skewness. The statistical analyses were conducted in RStudio v.4.1.2 with Student’s *t-*test, ANOVA, or a Kruskal–Wallis test. To investigate whether the relationship between *D. fangzhongdai* titer and root colonization rate among individual seedlings with leaf soft rot is linked to OMF identity, we used a generalized linear mixed model (GLMM) for negative binomial implemented in the *lmerTest* package (Kuznetsova *et al.,* 2017). Considering the possibility of different responses among individuals and infection occurrence, experimental batches and individual seedlings were used as a random factor and OMF identity and colonization rate were used as fixed factors. The final syntax was constructed without the interaction between OMF colonization rate and identity to produce, the most maximal yet parsimonious model: *glmer.nb* (colony-forming unit [CFU] count ∼ colonization rate + OMF identity + [1| experimental batch/treatment]). To determine the importance of OMF colonization rate and OMF identity in determining CFU count, the *model.sel* function of the *MuMIn* package (Bartoń, 2023) was applied to the constructed GLMM. The omitted factor with the largest change in the corrected Akaike information criterion value was regarded as the most important fixed factor.

## Results

### *Tulasnella calospora* and *S. vermifera* colonize *B. striata* roots with similar symbiotic affinity in protocorms

We established an *ex vitro* growing system for *B. striata* seedlings in a common potting mix and cultured OMF in liquid medium to obtain maximum mycelial growth and facilitate direct root inoculation. We tested the compatibility of *B. striata* with OMF candidates by direct root inoculation described in the current study, and by a symbiotic germination assay established previously (Fuji *et al*., 2020). Within 2 weeks, we confirmed the presence of pelotons in protocorms and roots, which is the primary sign of OM establishment (Figs. 1A, S2). In protocorms, OMF colonization started from the basal part and was restricted to the lower half until the formation of shoot primordium (Fig. S2), as reported earlier (Miura *et al*., 2019). In roots, OMF colonization occurred everywhere, except the quiescent center. The colonization rate was higher for *T. calospora* than for *S. vermifera* in protocorms (Fig. 1B) and in roots (Fig. 1C).

**Fig. 1.**
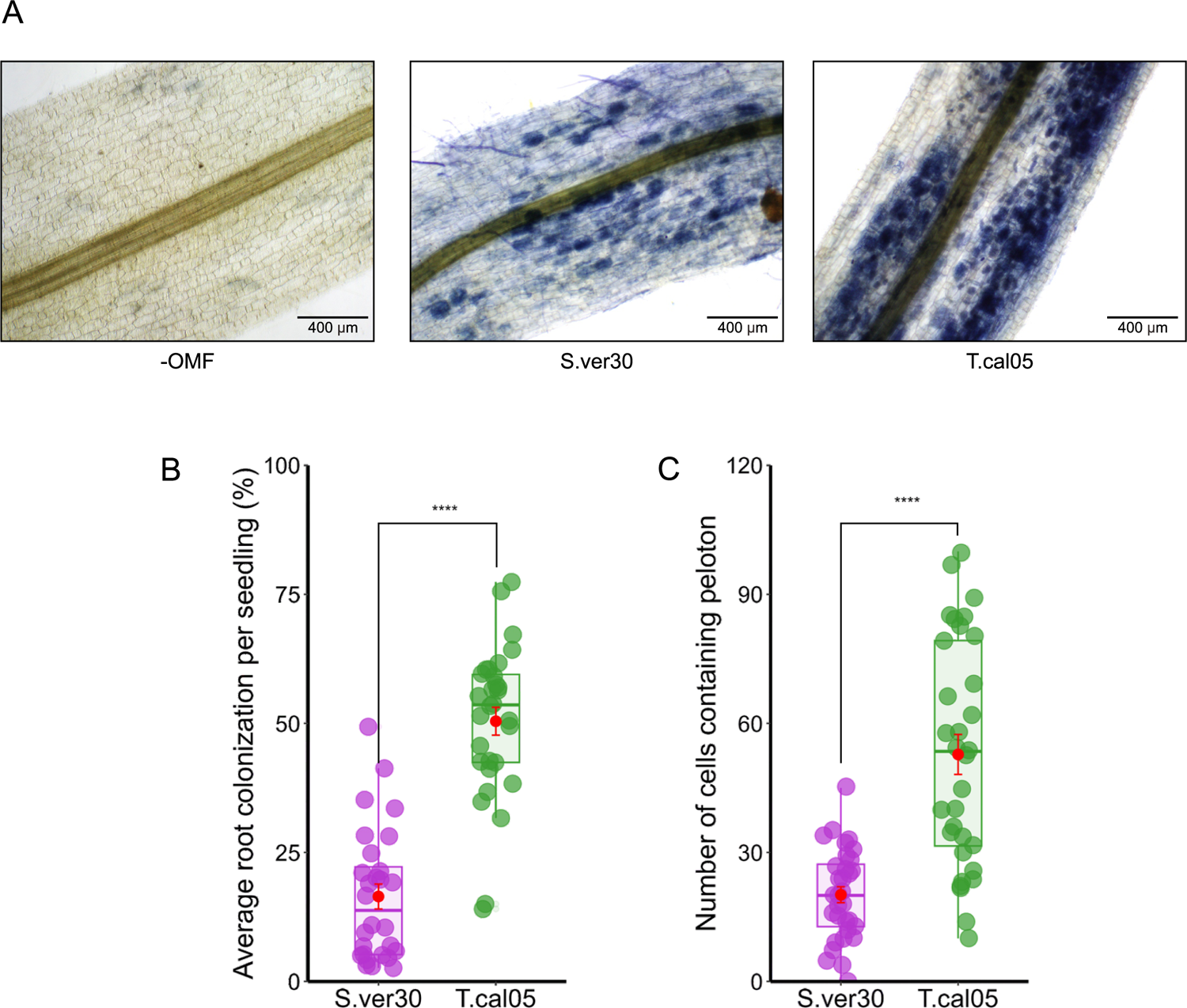
Colonization of *Bletilla striata* by orchid mycorrhizal fungi (OMF). (A) Root colonization. Pelotons are visible inside the root cortex. Scale bars, 400 µm. (B) Average root colonization rate. (C) Number of symbiotic cells per protocorm \*\*\*\**p* < 0.0001 (Student’s *t*-test). S.ver30, *Serendipita vermifera*; T.cal05, *Tulasnella calospora*.

### Erwinia chrysanthemi MAFF311045 is synonymous to Dickeya fangzhongdai

Our pathosystem for *B. striata* was based on a study on orchid infection by *Erwinia* species (Wu *et al*., 2011; Ye *et al*., 2019). We chose *Erwinia chrysanthemi* because of its ability to infect orchids. The *Erwinia* genus is rather complex (Samson *et al*., 2005). Our phylogenetic tree based on 16S ribosomal RNA sequences of various species within Pectobacteriaceae assigned the *E. chrysanthemi* MAFF311045 to *Dickeya fangzhongdai* (Fig. S3). Leaf inoculation with this pathogen resulted in a lesion and bacterial spread throughout the leaf, often causing the leaf to drop, in *B. striata*, *A. thaliana*, and *N. benthamiana* (Fig. S4).

### Induced systemic resistance is linked to the degree of OMF colonization

The effects of OMF colonization on ISR against pathogens were examined in the *B. striata*–*D. fangzhongdai* pathosystem. Both *S. vermifera* and *T. calospora* were able to colonize *B. striata* roots (Fig. 1). Upon *D. fangzhongdai* infection, lesions on detached leaves were wider in the presence of OMF colonization than in its absence (Fig. 2A). *Tulasnella calospora* colonization induced the defense response, as indicated by a reduced *D. fangzhongdai* titer (Fig. 2B).

**Fig. 2.**
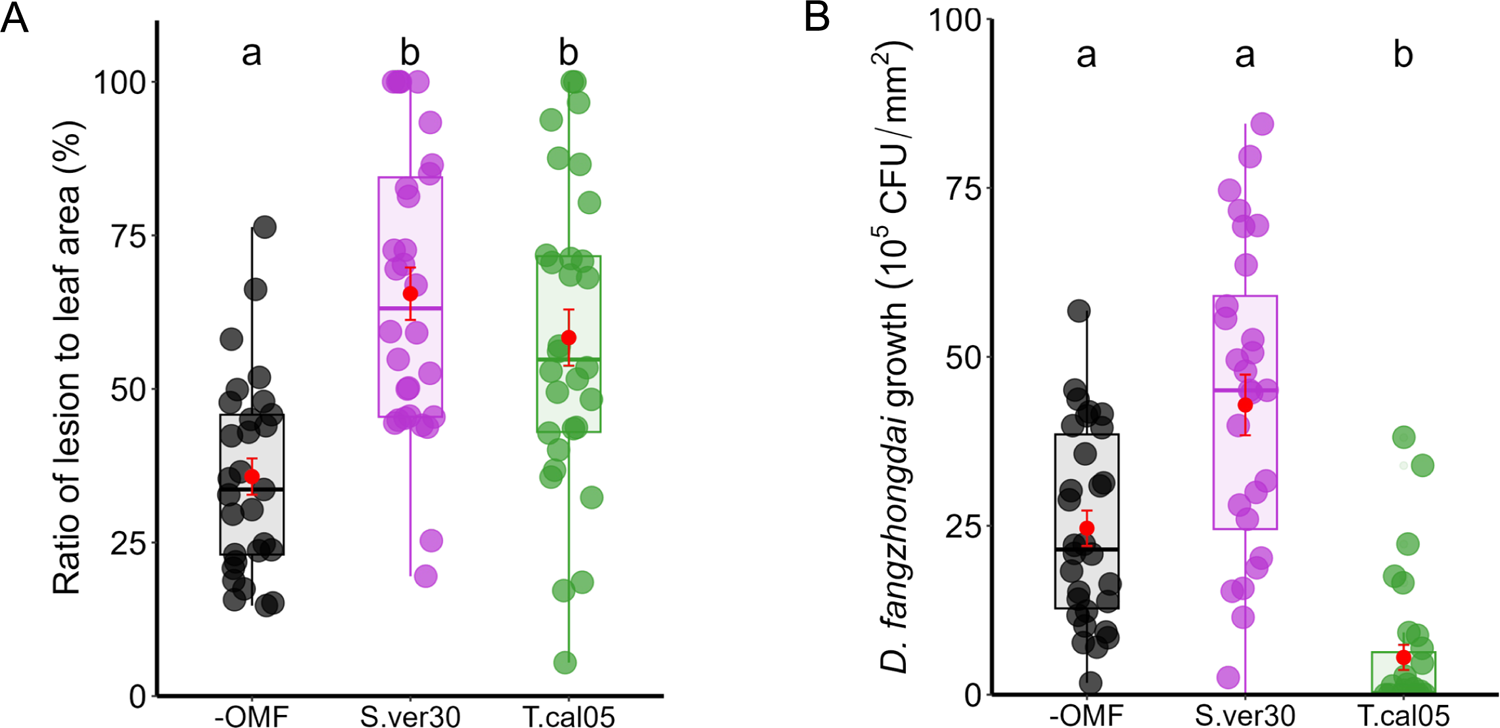
Induced systemic resistance on leaves upon *Dickeya fangzhongdai* infection. (A) Ratio of lesion to leaf area. (B) *D. fangzhongdai* colony-forming unit (CFU) count in leaves 3 days after infection (*p* < 0.05, Kruskal–Wallis test).

To further test the physiological changes associated with the induction of disease resistance, we assessed the indicators of photosynthetic damage and peroxide production. Photosynthetic damage is a general proxy for disease resistance in leaves, and reactive oxygen species are general inducers of disease resistance (Lu and Yao, 2018; Li and Kim, 2022). Regardless of lesion size, the necrotic area was relatively constant in all treatments, as revealed by trypan blue staining (Fig. 3A, B). The photosynthetic quantum yield was reduced in all infected leaves, but the photosynthetic damage caused by the infection was significantly smaller when *T. calospora* colonized the roots (Fig. C). Similarly, the decrease in total chlorophyll content of seedlings caused by *D. fangzhongdai* infection was lowest in the presence of *T. calospora* (Fig. 3D). Hydrogen peroxide was also accumulated in infected leaves (Fig. S5A), and its content was significantly higher in OMF-colonized seedlings (Fig. S5B). Overall, the ability of OMF to systemically induce disease resistance in orchids is not necessarily related to symptom appearance but rather to reduced pathogen proliferation. This is also supported by the data that both OMF identity and colonization rate strongly affected ISR (*n* = 88 plants; *p* < 0.001; Tables 1 and 2, Fig. S6).

**Fig. 3.**
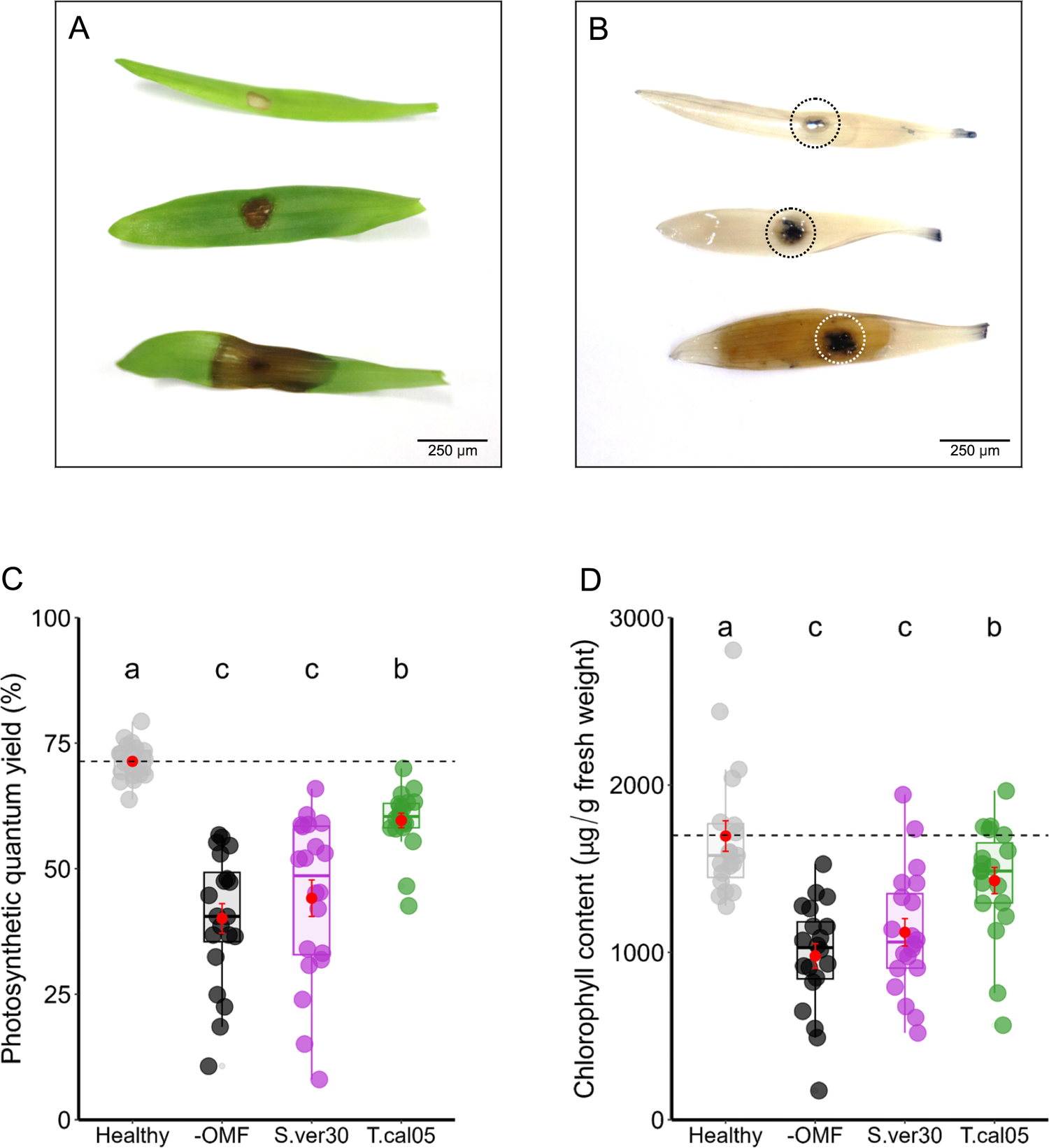
Leaf damage caused by *Dickeya fangzhongdai* infection. (A) Variability of lesion areas on infected leaves. (B) Necrotic areas (dotted circles) and lesions (brown) Leaves in (B) were stained with trypan blue. Scale bars, 250 µm. (C) Photosynthetic quantum yield and (D) total chlorophyll contents of infected leaves 3 days after infection and in healthy uninoculated seedlings (*p* < 0.05, Kruskal–Wallis test).

**Table 1.** Results of generalized linear mixed model.

| Fixed effects | Estimate | Standard error | z-value | p-value |
| --- | --- | --- | --- | --- |
| (Intercept) | 0.666351 | 0.215476 | 3.092 | < **0.01** |
| Colonization rate | -0.020733 | 0.004376 | -4.738 | < **0.001** |
| OMF identity | 0.035266 | 0.00508 | 6.943 | < **0.001** |

\*Significances are highlighted in bold

**Table 2.** Results of model comparison using corrected Akaike information criterion.

| Model | log likelihood | AICc | delta |
| --- | --- | --- | --- |
| Initial | -156.618 | 326.3 | 0 |
| Without colonization rate | -170.078 | 350.9 | 24.62 |
| Without OMF identity | -181.193 | 373.1 | 46.84 |

### Transcriptome analysis of leaves of *B. striata* seedlings inoculated with OMF and pathogen

The lowest *D. fangzhongdai* titer and photosynthetic damage in *T. calospora–*infected leaves led us to hypothesize that *T. calospora* colonization enhances disease resistance by systemically inducing changes in gene transcription. We analyzed transcriptomes of leaves upon root inoculation with *T. calospora* (T treatment), upon leaf infection by *D. fangzhongdai* (P treatment), or both (PT treatment).

Up- and downregulated DEGs were identified by comparisons among the three treatments (Tables S2*–*S4). A heatmap of DEG clustering showed distinct expression patterns among the treatments (Fig. 4A). A total of 2660 genes in P, 1391 in T, and 1628 in PT seedlings were upregulated, and 2987 in P, 2319 in T, and 857 in PT seedlings were downregulated. Among them, 6 DEGs (4 upregulated and 2 downregulated) were shared among all treatments, 1641 DEGs (397 upregulated and 1244 downregulated) between P and T, 63 DEGs (45 upregulated and 18 downregulated) between T and PT, and 5 upregulated DEGs between P and PT (Fig. 4A). The transcriptional profile was less similar between T and PT seedlings than between P and PT seedlings, as indicated by MDS plot (Fig. S7). These data demonstrated that OM symbiosis greatly affects the leaf transcriptome. Since both T and PT treatments involved *T. calospora* colonization, it systemically and conspicuously altered leaf transcriptional changes before and after *D. fangzhongdai* infection.

**Fig. 4.**
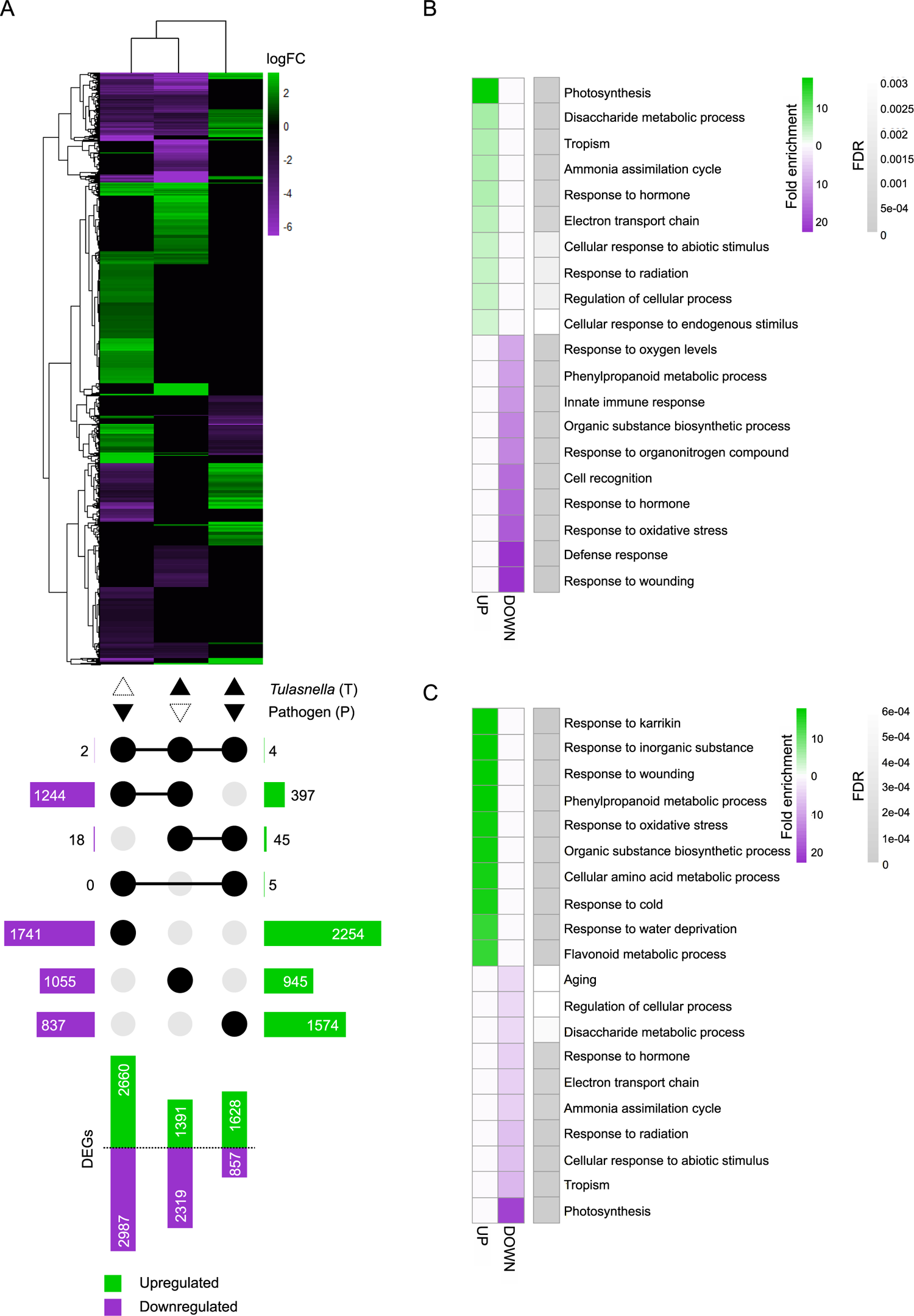
RNA-seq analysis. (A) Heatmap showing fold changes of differentially expressed genes (DEGs) in leaves of seedlings treated with the pathogen *Dickeya fangzhongdai* only (P); *Tulasnella calospora* only (T); or both (PT). An UpSet plot below the heatmap shows the number of upregulated and downregulated DEGs. (B) and (C) Heatmaps of the top 10 enriched biological process gene ontology (GO) terms for upregulated and downregulated DEGs in (B) T treatment and (C) PT treatment. FDR, false discovery rate. All GO terms are listed in Tables S5 and S6.

GO enrichment analysis of biological processes revealed that terms related to photosynthesis were overrepresented among the upregulated DEGs in T-treated seedling leaves. The categories of disaccharide metabolic process, tropism, ammonia assimilation cycle, and innate immune response were also enriched. Among downregulated DEGs, response to wounding, defense response, and response to oxidative stress were among the overrepresented terms (Fig. 4B, Table S5). In PT-treated seedlings, terms related to responses to karrikin, inorganic substance, and wounding were overrepresented among the upregulated DEGs, whereas photosynthesis, tropism, and ammonia assimilation cycle were among the overrepresented terms among downregulated DEGs (Fig. 4C, Table S6). These results implied that priming occurred upon *T. calospora* colonization by increasing primary metabolism and then resistance was increased upon pathogen infection.

### Comparisons of expression levels of phosphate and jasmonate/ethylene metabolism-related genes in rice and *B. striata*

We hypothesized that transcriptional patterns are conserved across different mycorrhizal types, especially in comparison with those in AM plants. To test this hypothesis, we compared our data with the published data on leaf transcriptional changes in rice (*Oryza sativa* ssp. *japonica* cv. ‘Loto’) inoculated with AM fungus *Funneliformis mosseae* (Campo and San Segundo, 2020). The most noticeable transcriptional changes in rice leaves were in phosphate- and jasmonate/ethylene metabolism-related genes. Jasmonate and ethylene are involved in resistance against necrotrophic pathogens and in ISR (Pieterse *et al.,* 2014). To identify the orthologous genes by SonicParanoid, we used rice genes as queries against *B. striata de novo* assembly (Table S7). We also used local TBLASTX (Table S8). The expression of the identified *B. striata* orthologs of phosphate metabolism– related genes did not necessarily mirror changes in the expression of rice genes (Fig. 5A, Table S8), but the identified genes for jasmonate and ethylene biosynthesis were similarly downregulated in rice and *B. striata* (Fig. 5B, Table S8). The expression of these genes was suppressed in T and P seedlings, but increased in PT seedlings.

**Fig. 5.**
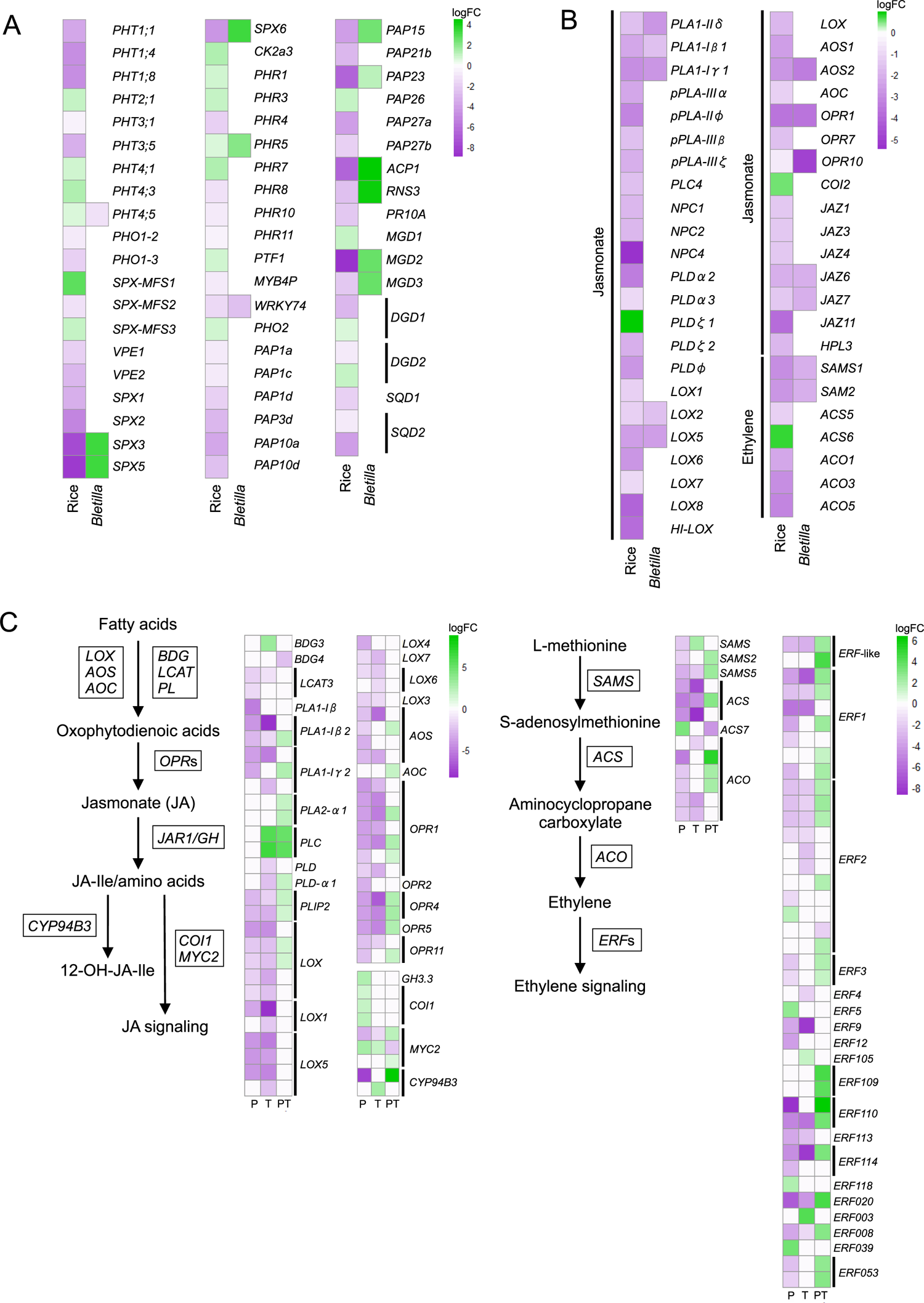
Comparative analysis of the expression patterns of phosphate-acquisition and defense hormone–related genes in leaves of rice (*Oryza sativa* spp. *japonica*, cv. Loto; Campo and Segundo, 2020) and *Bletilla striata* seedlings treated with the pathogen *Dickeya fangzhongdai* only (P); *Tulasnella calospora* only (T); or both (PT). (A) Phosphate metabolism and (B) jasmonate and ethylene signaling/metabolism. Orthologous genes of *B. striata* were identified by SonicParanoid and TBLASTX. (C) The expression of jasmonate- and ethylene-related genes of *B. striata* determined from transcriptomic analysis. See Tables S8 and S9 for details.

**Fig. 6.**
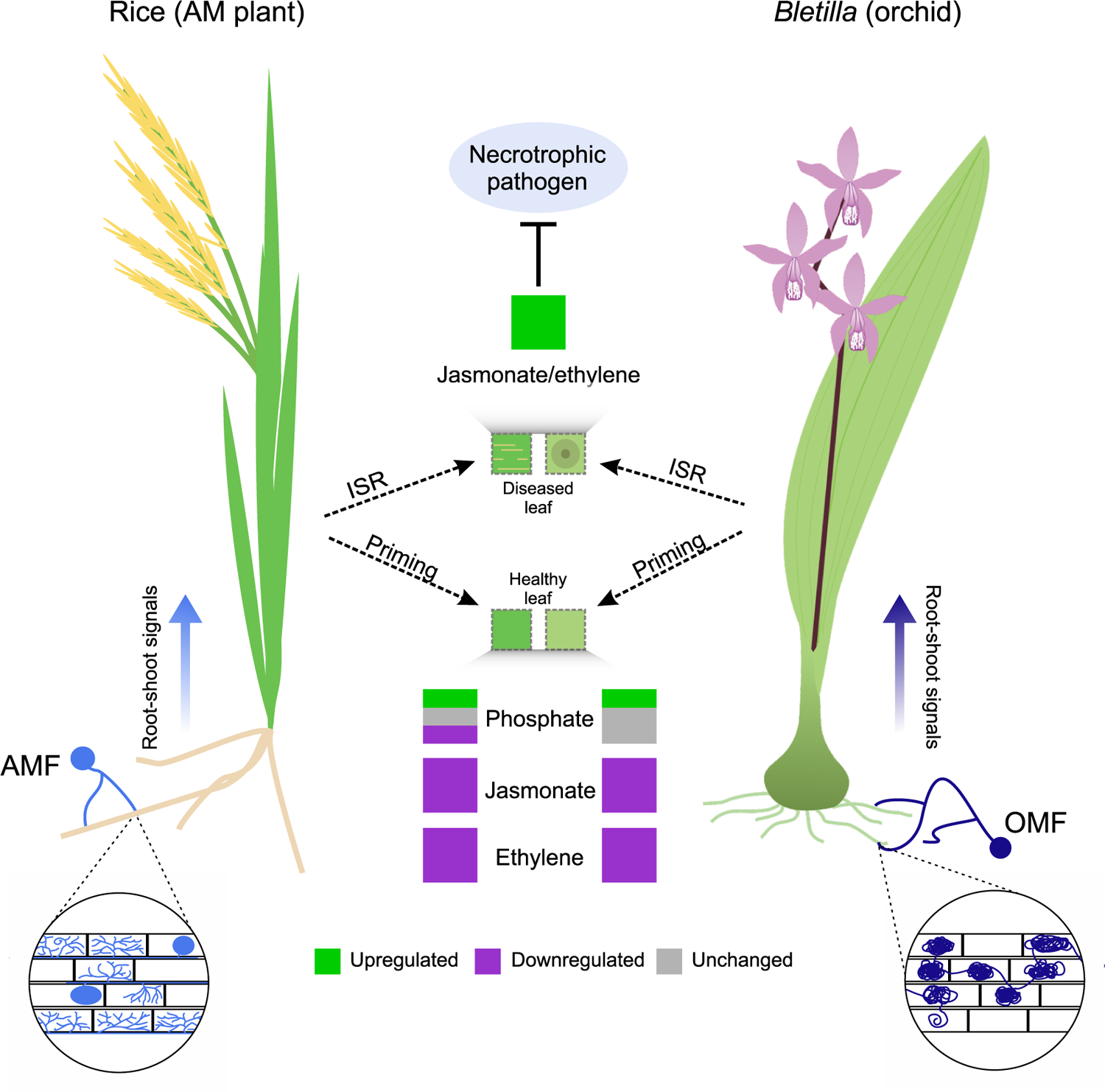
Commonalities in induced systemic resistance (ISR) primed by mycorrhizal fungus colonization in an AM plant (rice) and orchid (*Bletilla striata*). Despite structural differences, colonization by AMF and OMF causes priming by regulating the expression of genes related to primary metabolism (e.g., phosphate metabolism). This is more variable in AM plants than in orchids. However, the expression of genes related to the jasmonate and ethylene signaling pathways is downregulated in both species. When a necrotrophic pathogen infects the plants, AMF or OMF colonization triggers ISR in leaves, which increases the expression of genes involved in jasmonate and ethylene signaling pathways and in turn inhibits pathogen proliferation.

We further checked the expression patterns of homologous jasmonate and ethylene biosynthesis–related genes extracted from DEG lists (FDR < 0.05) and obtained similar results. The overall results showed that priming suppresses the jasmonate and ethylene signaling pathways and that leaf infection by pathogen activates them, confirming the occurrence of ISR in *B. striata* (Fig. 5C, Table S9).

## Discussion

In this study, we demonstrated that a compatible OMF colonization, *T. calospora,* systemically induces resistance against a necrotrophic pathogen as well as its possible role in defense priming in orchids. We also showed partial commonalities between AM and OM plants in aboveground transcriptomic changes during root colonization.

Mycorrhizal fungi improve host fitness, which varies among AM fungus–plant pairings (Säle *et al*., 2021; Cope *et al*., 2022; Guigard *et al*., 2023). The driving factors for the functional diversity are largely unknown, but plants may simply choose mycorrhizal fungi that provide the highest growth with minimal metabolic costs, which is important for survival. The same applies to orchids, which underwent rapid diversification in their growth habits during the Cenozoic (Dearnaley *et al*., 2012; Chomicki *et al*., 2015; Xing *et al*., 2019). Orchids with the highest demand for mycorrhizal association during early seed germination prioritize different OMF species as their partners (Fuji *et al.,* 2020; Otero *et al*., 2005; Meng *et al*., 2019; Pujasatria *et al*., 2022). Previously, we showed that *B. striata* associates with multiple strains of *Tulasnella* during seed germination (Fuji *et al*., 2020). It could be assumed that *B. striata* remains compatible with an OMF from seed germination until seedling development and quantified OMF colonization in seedlings with well-developed roots. Colonization during seed germination and root colonization was higher for *T. calospora* than for *S. vermifera.* Other *Bletilla* sister taxa, the members of tribe Arethuseae (e.g., *Arundina, Pleione*, and *Coelogyne*), are also associated with *Tulasnella*, as suggested by metagenomic studies and *in vitro* assays (Sathiyadash *et al*., 2014; Meng *et al*., 2019; Qin *et al*., 2019). These suggest that the convergent evolution of OM still involves mycorrhizal partner selection, in addition to molecular-level mechanisms shared with AM plants, such as the roles of phytohormones and transcriptional regulations (Miura *et al*., 2018, 2024).

*Dickeya fangzhongdai*, as well as *Erwinia sensu lato*, infects orchids (Fu *et al*., 2012; Alič *et al*., 2019; Ye *et al*., 2019; Zhou *et al*., 2021), *Arabidopsis* (Kraepiel *et al*., 2011), and tobacco (Sobiczewski *et al*., 2017), resulting in brownish soft necrotic lesions. The lesions, however, should not be used as the sole proxy of resistance; pathogen titer should be added as another criterion, as shown in this study. In *Phalaenopsis*, lesions are smaller upon OMF colonization (Wu *et al*., 2011). In this study on *B. striata*, the lesions were visually larger upon priming, but *D. fangzhongdai* titer was significantly lower. Whether these phenomena were due to the physiological difference between these orchid species is unknown. In this study, disease resistance against necrotrophic pathogen was systemically induced in orchids colonized by mycorrhizal fungi.

We investigated the potential similarity in ISR between orchids and AM plants through comparative transcriptomics. Upon root colonization by mycorrhizal fungi, the host plants alter various metabolic pathways (Fontana *et al*., 2009; Jacott *et al*., 2017; Goddard *et al*., 2021; Orine *et al*., 2022; Li *et al*., 2023); host plants increase their primary metabolic activity (nutrient uptake and remobilization) and temporarily lower the expression of defense-related genes (Schoenherr *et al*., 2019; Jing *et al*., 2022). In AM plants such as rice, genes associated with phosphate metabolism and phospholipid biosynthesis are upregulated, whereas non-phosphate lipids, which are later used in jasmonate biosynthesis, are downregulated (Campo and San Segundo, 2020). In *Bletilla*, the annotations related to phosphate metabolism are insufficient to conclude whether phosphate metabolism is also aligned with that of AM plants. Although it is unknown whether *Bletilla* does not prioritize phosphate metabolism or its phosphate-related genes are not differentially expressed, most studies on nutrient acquisition through OM show that orchids primarily focus on carbon and nitrogen from OMF (Stöckel *et al*., 2014; Fochi *et al*., 2016; Dearnaley and Cameron, 2017; Zahn *et al*., 2023). Inorganic phosphate is also taken up through OMF hyphae (Cameron *et al*., 2007; Davis *et al*., 2022) and its content is determined by trophic mode (Minasiewicz *et al*., 2023): mycoheterotrophic species have the highest phosphorus content, followed by partial mycoheterotrophic and autotrophic ones. Assuming that *B. striata* seedlings used in this study were already fully autotrophic, it is possible that most phosphate metabolism–related genes were not dominantly expressed even after plant colonization by OMF, unlike in AM plants.

On the other hand, after infection, the defense response was stronger after priming than before it. This is often indicated by an increase in defense-related gene expression and content of phytohormones, notably jasmonate and ethylene, which are involved in ISR and play a major role in defense against necrotrophic pathogens (Shoresh *et al*., 2005; Wasternack and Feussner, 2018). Mycorrhiza-colonized plants typically prioritize growth and thus allocate less resources to defense responses, including jasmonate biosynthesis (Huot *et al*., 2014). Once the leaf is attacked by a pathogen, jasmonate is readily increased for defense (Pieterse *et al.,* 2012). Our comparative transcriptome analysis based on annotations for rice showed similar patterns in *B. striata*, which highlights the conservation of physiological changes in mycorrhizal plants upon mycorrhizal fungal colonization, at least in monocots. Notably, genes involved in jasmonate and ethylene biosynthesis seemed to be upregulated after *D. fangzhongdai* infection in primed plants. In turn, *D. fangzhongdai* titer was reduced, similar to that in other pathogen infections of AM plants. Our transcriptional evidence further supports the presence of defense priming in orchids.

In summary, our study demonstrated that the association between orchid and mycorrhizal fungi leads to ISR, thus providing a new clue for commonalities between AM and OM. As in AM plants, OMF compatibility also reflects the best colonization. The presence of a particular OMF in a substrate is a key factor in orchid seedling establishment (Gowland *et al*., 2013; Pujasatria *et al*., 2022) to provide the protocorms with nutrients and to protect against stress. We also propose that the ISR in *B. striata* involves a complex mechanism in which regulation of metabolites, defense responses, photosynthesis, and oxidative stress responses is synchronized to reduce *D. fangzhongdai* proliferation during leaf soft rot onset. Most AM studies have focused on leaves (Fujita *et al*., 2022; Wang *et al*., 2022b), but priming could be more variable in orchids than in other plants because orchid diseases also occur on pseudobulbs, rhizomes, and even flowers (Ito and Aragaki, 1977; Swett and Uchida, 2015; Li *et al*., 2022). Thus, the priming effects in orchids are still largely unknown if one takes into account pathogen trophic modes, latency, and the attacked organs. Further studies are needed to clarify how the defense responses were activated at the molecular level, allowing to reveal more commonalities among AM plants and orchids as well as various types of pathogens.

## Acknowledgements

This work was supported by the National Institute for Basic Biology (NIBB) Cooperative Research Programs (Next-generation DNA Sequencing Initiative: 22NIBB403, 23NIBB401). We are grateful to the Japanese Ministry of Education, Culture, Sports, Science, and Technology (MEXT) for a scholarship to G.C.P., and to the Japan Society for the Promotion of Science (JSPS) for Research Fellowships for Young Scientists to C.M. We also thank the Data Integration and Analysis Facility, NIBB for supporting the RNA sequencing and providing computational resources.

## Author contributions

G.C.P., C.M., and H.K. designed the research; G.C.P. performed the experiments with the assistance of C.M. and H.K.; G.C.P., C.M., K.Y., and S.S. designed and performed the bioinformatic analyses; G.C.P., C.M., and H.K. wrote the manuscript, with comments from all authors.

## Data availability

Nucleotide sequence data from the RNA-seq analysis in this study are deposited in the DNA Data Bank of Japan (DDBJ) BioProject under the accession number PRJDB16858. Correspondence and requests for materials should be addressed to H.K.

## Supplementary data

**Fig. S1** Experimental design used in this study.

**Fig. S2** *Bletilla striata* protocorms. (A) A developing protocorm containing (B) pelotons. Scale bars (A) 200 µm, (B) 100 µm.

**Fig. S3** Maximum likelihood phylogenetic tree based on 16S sequences of various species within Pectobacteriaceae, including *Erwinia chrysanthemi* MAFF311045. The tree was constructed using the Hasegawa–Kishino–Yano model with gamma with invariant distribution (HKY+G+I) and 1000 bootstrap replicates. *Lonsdalea quercina* was used as the outgroup. The analysis was conducted in MEGA10 and visualized in iTOL v.6 (https://itol.embl.de).

**Fig. S4** Soft rot symptoms observed in leaves of (A) *Bletilla striata*, (B) *Arabidopsis thaliana*, and (C) *Nicotiana benthamiana*.

**Fig. S5** Peroxide production in infected leaves. (A) Peroxide accumulation visualized by diaminobenzidine staining (brown area). Scale bar, 5 mm. (B) Peroxide content (*p* < 0.05, Kruskal–Wallis test).

**Fig. S6** Scatter plot showing the two-dimensional relationship between OMF colonization rate and *Dickeya fangzhongdai* CFU count.

**Fig. S7** MDS plot showing separation of DEGs among all treatments.

**Table S1** Summary of RNA-seq analysis

**Table S2** Genes differentially expressed in infected uncolonized seedlings (P treatment)

**Table S3** Genes differentially expressed in uninfected *Tulasnella calospora*–colonized seedlings (T treatment)

**Table S4** Genes differentially expressed in infected *Tulasnella calospora*–colonized seedlings (PT treatment)

**Table S5** Regulated GO terms before infection in *Tulasnella calospora*–colonized seedlings (T treatment)

**Table S6** Regulated GO terms after infection in *Tulasnella calospora*–colonized seedlings (PT treatment)

**Table S7** *Bletilla*–rice orthologous genes identified by SonicParanoid

**Table S8** *Bletilla*–rice orthologous genes identified by TBLASTX

**Table S9** Jasmonate- and ethylene-related genes expressed in *Bletilla striata* used in Fig. 5C

## References

Alexa A, Rahnenfuhrer J. 2023. topGO: Enrichment Analysis for Gene Ontology.

Alič Š, Pédron J, Dreo T, Van Gijsegem F. 2019. Genomic characterisation of the new *Dickeya fangzhongdai* species regrouping plant pathogens and environmental isolates. BMC Genomics 20, 1–18.

Bidartondo MI, Burghardt B, Gebauer G, Bruns TD, Read DJ. 2004 Changing partners in the dark: isotopic and molecular evidence of ectomycorrhizal liaisons between forest orchids and trees. Proceedings of the Royal Society of London. Series B: Biological Sciences 271, 1799–1806.

Brundrett MC, Tedersoo L. 2018. Evolutionary history of mycorrhizal symbioses and global host plant diversity. New Phytologist 220, 1108–1115.

Cameron DD, Johnson I, Leake JR, Read DJ. 2007. Mycorrhizal acquisition of inorganic phosphorus by the green-leaved terrestrial orchid *Goodyera repen*s. Annals of Botany 99, 831–834.

Cameron DD, Johnson I, Read DJ, Leake JR. 2008. Giving and receiving: Measuring the carbon cost of mycorrhizas in the green orchid, *Goodyera repens*. New Phytologist 180, 176–184.

Campo S, San Segundo B. 2020. Systemic induction of phosphatidylinositol-based signaling in leaves of arbuscular mycorrhizal rice plants. Scientific Reports 10, 1–17.

Cating RA, Palmateer AJ. 2011. Bacterial Soft Rot of *Oncidium* Orchids Caused by a *Dickeya* sp. (Pectobacterium chrysanthemi) in Florida. Plant Disease 95, 74.

Chen S, Zhou Y, Chen Y, Gu J, Gu Z, Eils R, Schlesner M. 2018. Fastp: An ultra-fast all-in-one FASTQ preprocessor. Bioinformatics 34, 2847–2849.

Chomicki G, Bidel LPR, Ming F, Coiro M, Zhang X, Wang Y, Baissac Y, Jay-Allemand C, Renner SS. 2015. The velamen protects photosynthetic orchid roots against UV-B damage, and a large dated phylogeny implies multiple gains and losses of this function during the Cenozoic. New Phytologist 205, 1330–1341.

Cope KR, Kafle A, Yakha JK, Pfeffer PE, Strahan GD, Garcia K, Subramanian S, Bücking H. 2022. Physiological and transcriptomic response of *Medicago truncatula* to colonization by high- or low-benefit arbuscular mycorrhizal fungi. Mycorrhiza 32, 281–303.

Cosentino S, Iwasaki W. 2019. SonicParanoid: Fast, accurate and easy orthology inference. Bioinformatics 35, 149–151.

David L, Harmon AC, Chen S. 2019. Plant immune responses - from guard cells and local responses to systemic defense against bacterial pathogens. Plant Signaling and Behavior 14, 1–9.

Davis B, Lim WH, Lambers H, Dixon KW, Read DJ. 2022. Inorganic phosphorus nutrition in green-leaved terrestrial orchid seedlings. Annals of Botany 129, 669–678.

Dearnaley JDW, Cameron DD. 2017. Nitrogen transport in the orchid mycorrhizal symbiosis – further evidence for a mutualistic association. New Phytologist 213, 10– 12.

Dearnaley JDW, Martos F, Selosse MA. 2012. Orchid mycorrhizas: Molecular ecology, physiology, evolution and conservation aspects. In: Hock B, ed. Fungal Associations. Heidelberg: Springer Berlin, 207–230.

Dreischhoff S, Das IS, Jakobi M, Kasper K, Polle A. 2020. Local Responses and Systemic Induced Resistance Mediated by Ectomycorrhizal Fungi. Frontiers in Plant Science 11, 1–20.

Eck JL, Kytöviita MM, Laine AL. 2022. Arbuscular mycorrhizal fungi influence host infection during epidemics in a wild plant pathosystem. New Phytologist 236, 1922– 1935.

Fochi V, Chitarra W, Kohler A, et al. 2016. Fungal and plant gene expression in the *Tulasnella calospora - Serapias vomeracea* symbiosis provides clues about nitrogen pathways in orchid mycorrhizas. New Phytologist 213, 365–379.

Fogell DJ, Kundu S, Roberts DL. 2019. Genetic homogenisation of two major orchid viruses through global trade-based dispersal of their hosts. Plants People Planet 1, 356– 362.

Fontana A, Reichelt M, Hempel S, Gershenzon J, Unsicker SB. 2009. The effects of arbuscular mycorrhizal fungi on direct and indirect defense metabolites of *Plantago lanceolata* L. Journal of Chemical Ecology 35, 833–843.

Fracchia S, Aranda-Rickert A, Rothen C, Sede S. 2016. Associated fungi, symbiotic germination and in vitro seedling development of the rare Andean terrestrial orchid *Chloraea riojana*. Flora 224, 106–111.

Fu SF, Tsai TM, Chen YR, Liu CP, Haiso LJ, Syue LH, Yeh HH, Huang HJ. 2012. Characterization of the early response of the orchid, *Phalaenopsis amabilis*, to *Erwinia chrysanthemi* infection using expression profiling. Physiologia Plantarum 145, 406– 425.

Fuji M, Miura C, Yamamoto T, Komiyama S, Suetsugu K, Yagame T, Yamato M, Kaminaka H. 2020. Relative effectiveness of *Tulasnella* fungal strains in orchid mycorrhizal symbioses between germination and subsequent seedling growth. Symbiosis 81, 53–63.

Fujita M, Kusajima M, Fukagawa M, Okumura Y, Nakajima M, Akiyama K, Asami T, Yoneyama K, Kato H, Nakashita H. 2022. Response of tomatoes primed by mycorrhizal colonization to virulent and avirulent bacterial pathogens. Scientific Reports 12, 1–12.

Girlanda M, Segreto R, Cafasso D, Liebel HT, Rodda M, Ercole E, Cozzolino S, Gebauer G, Perotto S. 2011. Photosynthetic Mediterranean meadow orchids feature partial mycoheterotrophy and specific mycorrhizal associations. American Journal of Botany 98, 1148–1163.

Goddard ML, Belval L, Martin IR, Roth L, Laloue H, Deglène-Benbrahim L, Valat L, Bertsch C, Chong J. 2021. Arbuscular Mycorrhizal Symbiosis Triggers Major Changes in Primary Metabolism Together With Modification of Defense Responses and Signaling in Both Roots and Leaves of *Vitis vinifera*. Frontiers in Plant Science 12, 1–16.

Gowland KM, Van Der Merwe MM, Linde CC, Clements MA, Nicotra AB. 2013. The host bias of three epiphytic Aeridinae orchid species is reflected, but not explained, by mycorrhizal fungal associations. American Journal of Botany 100, 764–777.

Gu Z, Eils R, Schlesner M. 2016. Complex heatmaps reveal patterns and correlations in multidimensional genomic data. Bioinformatics 32, 2847–2849.

Guigard L, Jobert L, Busset N, Moulin L, Czernic P. 2023. Symbiotic compatibility between rice cultivars and arbuscular mycorrhizal fungi genotypes affects rice growth and mycorrhiza-induced resistance. Frontiers in Plant Science 14, 1–17.

Haney CH, Wiesmann CL, Shapiro LR, et al. 2018. Rhizosphere-associated *Pseudomonas* induce systemic resistance to herbivores at the cost of susceptibility to bacterial pathogens. Molecular Ecology 27, 1833–1847.

Hélias V, Hamon P, Huchet E, Wolf J V.D., Andrivon D. 2012. Two new effective semiselective crystal violet pectate media for isolation of *Pectobacterium* and *Dickeya*. Plant Pathology 61, 339–345.

Huot B, Yao J, Montgomery BL, He SY. 2014. Growth-defense tradeoffs in plants: A balancing act to optimize fitness. Molecular Plant 7, 1267–1287.

Ito JS, Aragaki M. 1977. *Botrytis* Blossom Blight of *Dendrobium*. Phytopathology 67, 820–824.

Jacott CN, Murray JD, Ridout CJ. 2017. Trade-offs in arbuscular mycorrhizal symbiosis: Disease resistance, growth responses and perspectives for crop breeding. Agronomy 7, 1–18.

Jing S, Li Y, Zhu L, Su J, Yang T, Liu B, Ma B, Ma F, Li M, Zhang M. 2022. Transcriptomics and metabolomics reveal effect of arbuscular mycorrhizal fungi on growth and development of apple plants. Frontiers in Plant Science 13, 1–16.

Joko T, Subandi A, Kusumandari N, Wibowo A, Priyatmojo A. 2014. Activities of plant cell wall-degrading enzymes by bacterial soft rot of orchid. Archives Of Phytopathology And Plant Protection 47, 1239–1250.

Kakouridis A, Hagen JA, Kan MP, Mambelli S, Feldman LJ, Herman DJ, Weber PK, Pett-Ridge J, Firestone MK. 2022. Routes to roots: direct evidence of water transport by arbuscular mycorrhizal fungi to host plants. New Phytologist 236, 210–221.

Kawahara Y, de la Bastide M, Hamilton JP, et al. 2013. Improvement of the *Oryza sativa* Nipponbare reference genome using next generation sequence and optical map data. Rice 6, 3–10.

Keith LM, Sewake KT, Zee FT. 2005. Isolation and characterization of *Burkholderia gladioli* from orchids in Hawaii. Plant Disease 89, 1273–1278.

Khamtham J, Akarapisan A. 2019. *Acidovorax avenae* subsp. *cattleyae* causes bacterial brown spot disease on terrestrial orchid *Habenaria lindleyana* in Thailand. Journal of Plant Pathology 101, 31–37.

Kraepiel Y, Pédron J, Patrit O, Simond-Côte E, Hermand V, van Gijsegem F. 2011. Analysis of the plant *Bos1* mutant highlights necrosis as an efficient defence mechanism during *D. dadantii/Arabidospis thaliana* interaction. PLoS ONE 6, 1–10.

Kuga Y, Sakamoto N, Yurimoto H. 2014. Stable isotope cellular imaging reveals that both live and degenerating fungal pelotons transfer carbon and nitrogen to orchid protocorms. New Phytologist 202, 594–605.

Kuznetsova A, Brockhoff PB, Christensen RHB. 2017. lmerTest Package: Tests in Linear Mixed Effects Models. Journal of Statistical Software 82, 1–26.

Langmead B, Salzberg SL. 2012. Fast gapped-read alignment with Bowtie 2. Nature Methods 9, 357–359.

Lee YA, Yu CP. 2006. A differential medium for the isolation and rapid identification of a plant soft rot pathogen, *Erwinia chrysanthemi*. Journal of Microbiological Methods 64, 200–206.

Li J, Zhang M, Yang Z, Li C. 2022. *Botrytis cinerea* causes flower gray mold in *Gastrodia elata* in China. Crop Protection 155, 1–4.

Li M, Kim C. 2022. Chloroplast ROS and stress signaling. Plant Communications 3, 1–15.

Li Y, Nan Z, Matthew C, Wang Y, Duan T. 2023. Arbuscular mycorrhizal fungus changes alfalfa (*Medicago sativa*) metabolites in response to leaf spot (*Phoma medicaginis*) infection, with subsequent effects on pea aphid (*Acyrthosiphon pisum*) behavior. New Phytologist 239, 286–300.

Liu J, Maldonado-Mendoza I, Lopez-Meyer M, Cheung F, Town CD, Harrison MJ. 2007. Arbuscular mycorrhizal symbiosis is accompanied by local and systemic alterations in gene expression and an increase in disease resistance in the shoots. Plant Journal 50, 529–544.

Lopes UP, Zambolim L, Pereira OL. 2009. First report of *Lasiodiplodia theobromae* causing leaf blight on the orchid *Catasetum fimbriatum* in Brazil. Australasian Plant Disease Notes 4, 64–65.

Lu Y, Yao J. 2018. Chloroplasts at the crossroad of photosynthesis, pathogen infection and plant defense. International Journal of Molecular Sciences 19, 1–37.

Marquez N, Giachero ML, Gallou A, Debat HJ, Cranenbrouck S, Di Rienzo JA, Pozo MJ, Ducasse DA, Declerck S. 2018. Transcriptional changes in mycorrhizal and nonmycorrhizal soybean plants upon infection with the fungal pathogen *Macrophomina phaseolina*. Molecular Plant-Microbe Interactions 31, 842–855.

McGonigle TP, Miller MH, Evans DG, Fairchild GL, Swan JA. 1990. A new method which gives an objective measure of colonization of roots by vesicular-arbuscular mycorrhizal fungi. New Phytologist 115, 495–501.

McKendrick SL, Leake JR, Read DJ. 2000. Symbiotic germination and development of myco-heterotrophic plants in nature: transfer of carbon from ectomycorrhizal *Salix repens* and *Betula pendula* to the orchid *Corallorhiza trifida* through shared hyphal connections. New Phytologist 145, 539–548.

Meng YY, Zhang WL, Selosse MA, Gao JY. 2019. Are fungi from adult orchid roots the best symbionts at germination? A case study. Mycorrhiza 29, 541–547.

Minasiewicz J, Zwolicki A, Figura T, Novotná A, Bocayuva MF, Jersáková J, Selosse MA. 2023. Stoichiometry of carbon, nitrogen and phosphorus is closely linked to trophic modes in orchids. BMC Plant Biology 23, 1–11.

Miura C, Furui Y, Yamamoto T, et al. 2024. Autoactivation of mycorrhizal symbiosis signaling through gibberellin deactivation in orchid seed germination. Plant Physiology 194, 1–18.

Miura C, Saisho M, Yagame T, Yamato M, Kaminaka H. 2019. *Bletilla striata* (Orchidaceae) seed coat restricts the invasion of fungal hyphae at the initial stage of fungal colonization. Plants 8, 1–11.

Miura C, Yamaguchi K, Miyahara R, Yamamoto T, Fuji M, Yagame T, Imaizumi-Anraku H, Yamato M, Shigenobu S, Kaminaka H. 2018. The Mycoheterotrophic Symbiosis Between Orchids and Mycorrhizal Fungi Possesses Major Components Shared with Mutualistic Plant-Mycorrhizal Symbioses. Molecular Plant Microbe Interactions 31, 1032–1047.

Orine D, Defossez E, Vergara F, Uthe H, van Dam NM, Rasmann S. 2022. Arbuscular mycorrhizal fungi prevent the negative effect of drought and modulate the growth-defence trade-off in tomato plants. Journal of Sustainable Agriculture and Environment 1, 177–190.

Otero JT, Bayman P, Ackerman JD. 2005. Variation in mycorrhizal performance in the epiphytic orchid *Tolumnia variegata* in vitro: The potential for natural selection. Evolutionary Ecology 19, 29–43.

Parniske M. 2008 Arbuscular mycorrhiza: the mother of plant root endosymbioses. Nature Reviews Microbiology 6, 763–775.

van Peer R, Niemann GJ, Schippers B. 1991. Induced Resistance and Phytoalexin Accumulation in Biological Control of Fusarium Wilt of Carnation by *Pseudomonas* sp. Strain WCS417r. Pythopathology 81, 728–734.

Pérez FJ, Rubio S. 2006. An improved chemiluminescence method for hydrogen peroxide determination in plant tissues. Plant Growth Regulation 48, 89–95.

Peterson RL, Massicote HB, Melville LH. 2004. Mycorrhizas: Anatomy and Cell Biology. Ottawa: NRC Research Press.

Pieterse CMJ, Van der Does D, Zamioudis C, Leon-Reyes A, Van Wees SCM. 2012. Hormonal modulation of plant immunity. Annual Review of Cell and Developmental Biology 28, 489–521.

Pieterse CMJ, Zamioudis C, Berendsen RL, Weller DM, Van Wees SCM, Bakker PAHM. 2014. Induced systemic resistance by beneficial microbes. Annual Review of Phytopathology 52, 347–375.

Pujasatria GC, Nishiguchi I, Miura C, Yamato M, Kaminaka H. 2022. Orchid mycorrhizal fungi and ascomycetous fungi in epiphytic *Vanda falcata* roots occupy different niches during growth and development. Mycorrhiza 32, 481–495.

Qin J, Zhang W, Ge ZW, Zhang SB. 2019. Molecular identifications uncover diverse fungal symbionts of *Pleione* (Orchidaceae). Fungal Ecology 37, 19–29.

Ross AF. 1966. Systemic effects of local lesion formation. In: Beemster ABR, Dijkstra J, eds. Viruses of Plants: Their Isolation, Purification, and Characterization: the Mechanism of Plant Virus Infection, Synthesis of Viral Protein and Viral Nucleic Acid, and Plant Reactions Evoked by Viruses. Wageningen: North-Holland Publishing Company, 127–150.

Säle V, Palenzuela J, Azcón-Aguilar C, Sánchez-Castro I, da Silva GA, Seitz B, Sieverding E, van der Heijden MGA, Oehl F. 2021. Ancient lineages of arbuscular mycorrhizal fungi provide little plant benefit. Mycorrhiza 31, 559–576.

Samson R, Legendre JB, Christen R, Fischer-Le Saux M, Achouak W, Gardan L. 2005. Transfer of Pectobacterium chrysanthemi (Burkholder et al. 1953) Brenner et al. 1973 and Brenneria paradisiaca to the genus Dickeya gen. nov. as Dickeya chrysanthemi comb. nov. and Dickeya paradisiaca comb. nov. and delineation of four novel species, Dick. International Journal of Systematic and Evolutionary Microbiology 55, 1415–1427.

Sathiyadash K, Muthukumar T, Murugan SB, Sathishkumar R, Pandey RR. 2014. In vitro symbiotic seed germination of South Indian endemic orchid *Coelogyne nervosa*. Mycoscience 55, 183–189.

Schoenherr AP, Rizzo E, Jackson N, Manosalva P, Gomez SK. 2019. Mycorrhiza-induced resistance in potato involves priming of defense responses against cabbage looper (Noctuidae: Lepidoptera). Environmental Entomology 48, 370–381.

Selosse MA, Minasiewicz J, Boullard B. 2017. An annotated translation of Noël Bernard’s 1899 article ‘On the germination of *Neottia nidus-avis*’. Mycorrhiza 27, 611–618.

Shoresh M, Yedidia I, Chet I. 2005. Involvement of jasmonic acid/ethylene signaling pathway in the systemic resistance induced in cucumber by *Trichoderma asperellum* T203. Phytopathology 95, 76–84.

Silva M, Pereira OL. 2007. First report of *Guignardia endophyllicola* leaf blight on *Cymbidium* (Orchidaceae) in Brazil. Australasian Plant Disease Notes 2, 31–32.

Smith SE, Read D. 2008. Mycorrhizal Symbiosis. New York: Academic Press Inc.

Smyth GK, Ritchie ME, Law CW, Alhamdoosh M, Su S, Dong X, Tian L. 2018. RNA-seq analysis is easy as 1-2-3 with limma, Glimma and edgeR. F1000Research 5, 1–30.

Sobiczewski P, Iakimova ET, Mikiciński A, Węgrzynowicz-Lesiak E, Dyki B. 2017. Necrotrophic behaviour of *Erwinia amylovora* in apple and tobacco leaf tissue. Plant Pathology 66, 842–855.

Srivastava S, Kadooka C, Uchida JY. 2018. *Fusarium* species as pathogen on orchids. Microbiological Research 207, 188–195.

Stöckel M, Těšitelová T, Jersáková J, Bidartondo MI, Gebauer G. 2014. Carbon and nitrogen gain during the growth of orchid seedlings in nature. New Phytologist 202, 606–615.

Strullu-Derrien C, Selosse MA, Kenrick P, Martin FM. 2018. The origin and evolution of mycorrhizal symbioses: from palaeomycology to phylogenomics. New Phytologist 220, 1012–1030.

Suharjo R, Sawada H, Takikawa Y. 2014. Phylogenetic study of Japanese *Dickeya* spp. and development of new rapid identification methods using PCR-RFLP. Journal of General Plant Pathology 80, 237–254.

Suwannarach N, Kumla J, Lumyong S. 2018. Leaf spot on *Cattleya* orchid caused by *Neoscytalidium orchidacearum* in Thailand. Canadian Journal of Plant Pathology 40, 109–114.

Swett CS, Uchida JY. 2015. Characterization of *Fusarium* diseases on commercially grown orchids in Hawaii. Plant Pathology 64, 648–654.

Tsai CF, Huang CH, Wu FH, Lin CH, Lee CH, Yu SS, Chan YK, Jan FJ. 2022. Intelligent image analysis recognizes important orchid viral diseases. Frontiers in Plant Science 13, 1–20.

Umata H, Ota Y, Yamada M, Watanabe Y, Gale SW. 2013. Germination of the fully myco-heterotrophic orchid *Cyrtosia septentrionalis* is characterized by low fungal specificity and does not require direct seed-mycobiont contact. Mycoscience 54, 343– 352.

Veldre V, Abarenkov K, Bahram M, Martos F, Selosse MA, Tamm H, Kõljalg U, Tedersoo L. 2013. Evolution of nutritional modes of Ceratobasidiaceae (Cantharellales, Basidiomycota) as revealed from publicly available ITS sequences. Fungal Ecology 6, 256–268.

Vierheilig H, Coughlan AP, Wyss U, Piche Y. 1998. Ink and Vinegar, a Simple Staining Technique for Arbuscular-Mycorrhizal Fungi. Appl Environ Microbiol 64, 5004–5007.

Vlot AC, Sales JH, Lenk M, Bauer K, Brambilla A, Sommer A, Chen Y, Wenig M, Nayem S. 2021. Systemic propagation of immunity in plants. New Phytologist 229, 1234–1250.

Wang H, Hao Z, Zhang X, Xie W, Chen B. 2022a. Arbuscular mycorrhizal fungi induced plant resistance against *Fusarium* wilt in jasmonate biosynthesis defective mutant and wild type of tomato. Journal of Fungi 8, 1–14.

Wang M, Tang W, Xiang L, Chen X, Shen X, Yin C, Mao Z. 2022b. Involvement of *MdWRKY40* in the defense of mycorrhizal apple against *Fusarium solani*. BMC Plant Biology 22, 1–15.

Wasternack C, Feussner I. 2018. The Oxylipin Pathways: Biochemistry and Function. Annual Review of Plant Biology 69, 363–386.

Wei G, Kloepper JW, Tuzun S. 1991. Induction of Systemic Resistance of Cucumber to *Colletotrichum orbiculare* by Select Strains of Plant Growth-Promoting Rhizobacteria. Phytopathology 81, 1508–1512.

Wei XY, Deng WL, Chu CC. 2021. Phylogenetic and phenotypic analyses on *Dickeya* spp. isolated from different host plants in Taiwan. Journal of Phytopathology 169, 678– 691.

Wellburn AR. 1994. The Spectral Determination of Chlorophylls a and b, as well as Total Carotenoids, Using Various Solvents with Spectrophotometers of Different Resolution. Journal of Plant Physiology 144, 307–313.

Wu PH, Huang DD, Chang DCN. 2011. Mycorrhizal symbiosis enhances *Phalaenopsis* orchid’s growth and resistance to *Erwinia chrysanthemi*. African Journal of Biotechnology 10, 10095–10100.

Xing X, Jacquemyn H, Gai X, Gao Y, Liu Q, Zhao Z, Guo S. 2019. The impact of life form on the architecture of orchid mycorrhizal networks in tropical forest. Oikos 128, 1254–1264.

Yamamoto T, Miura C, Fuji M, Nagata S, Otani Y, Yagame T, Yamato M, Kaminaka H. 2017. Quantitative evaluation of protocorm growth and fungal colonization in *Bletilla striata* (Orchidaceae) reveals less-productive symbiosis with a non-native symbiotic fungus. BMC Plant Biology 17, 1–10.

Ye W, Jiang J, Lin Y, Yeh KW, Lai Z, Xu X, Oelmüller R. 2019. Colonisation of *Oncidium* orchid roots by the endophyte *Piriformospora indica* restricts *Erwinia chrysanthemi* infection, stimulates accumulation of NBS-LRR resistance gene transcripts and represses their targeting micro-RNAs in leaves. BMC Plant Biology 19, 1–16.

Yeh CM, Chung KM, Liang CK, Tsai WC. 2019. New insights into the symbiotic relationship between orchids and fungi. Applied Sciences 9, 1–14.

Zahn FE, Söll E, Chapin TK, Wang D, Gomes SIF, Hynson NA, Pausch J, Gebauer G. 2023. Novel insights into orchid mycorrhiza functioning from stable isotope signatures of fungal pelotons. New Phytologist 239, 1449–1463.

Zhou A, Nie J, Tian Y, Chuan J, Hu B, Zou J, Li X. 2021. First Report of Dickeya fangzhongdai Causing Soft Rot in Orchid in Canada. Plant Disease 105, 4149.

