## Supplemental Figure 1-7 for "Colonization by orchid mycorrhizal fungi primes induced systemic resistance against a necrotrophic pathogen"

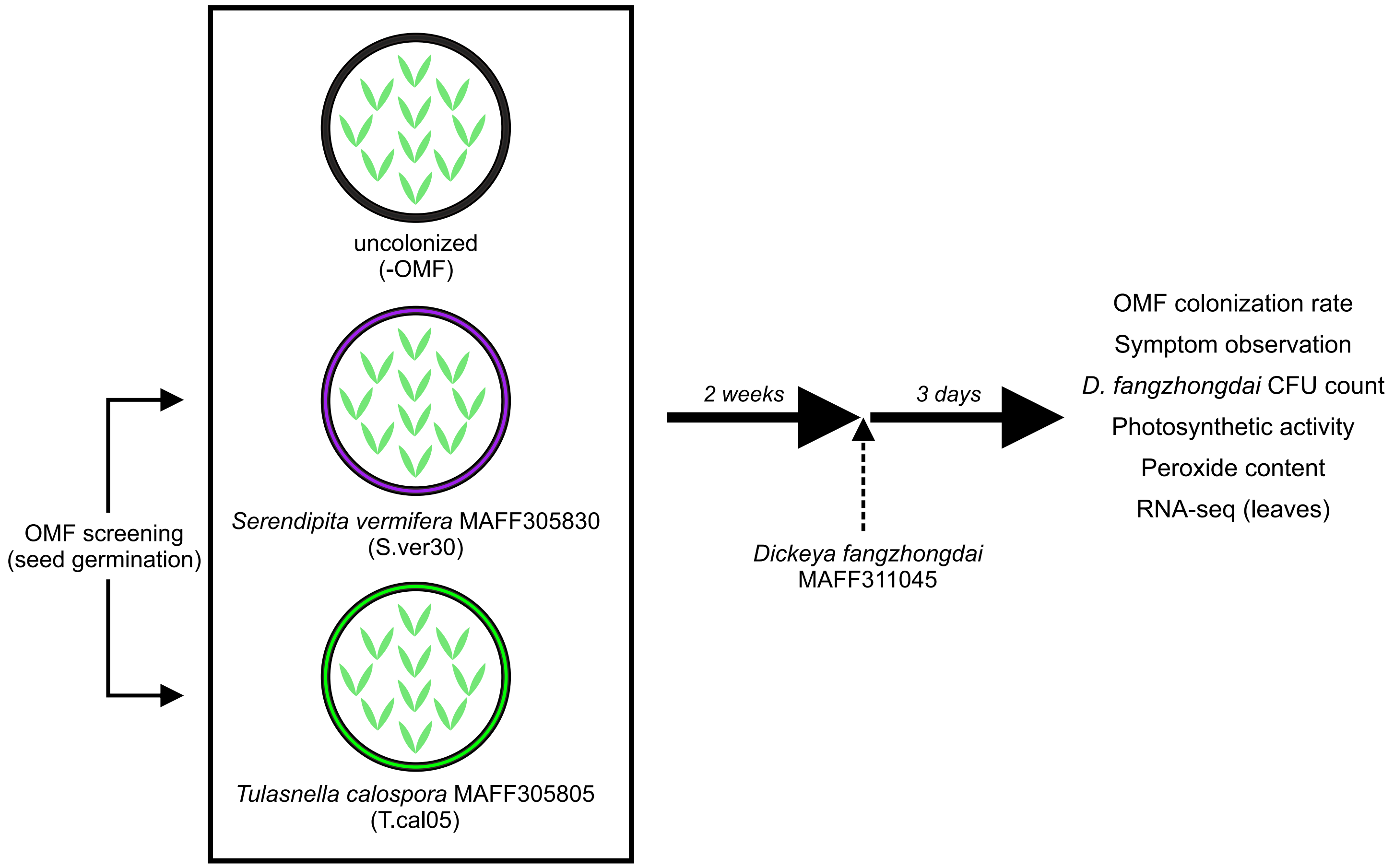


**Fig. S1** Experimental design in this study.

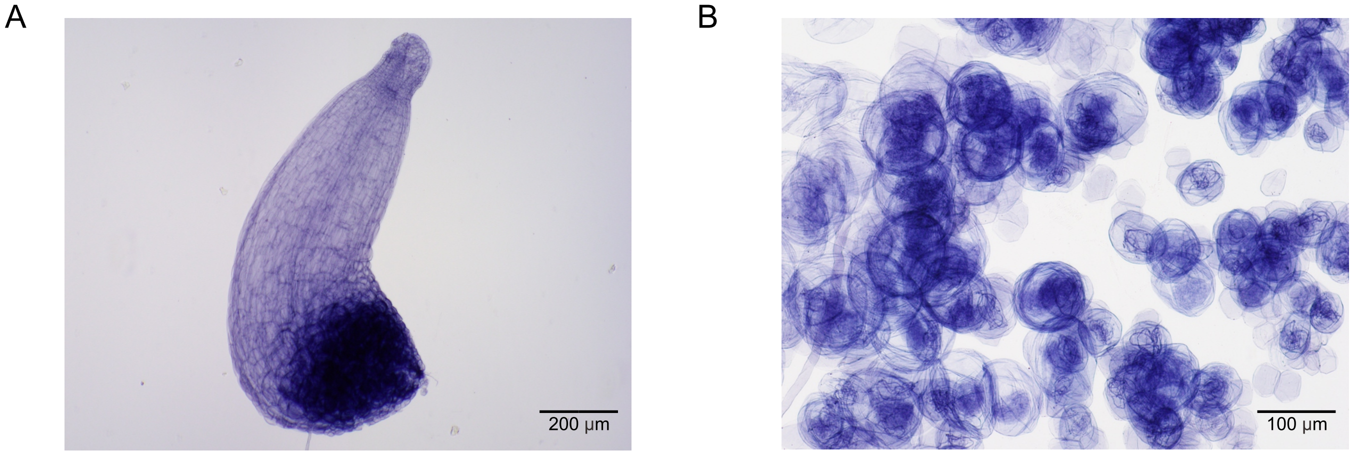


**Fig. S2** *Bletilla striata* protocorms. (A) A developing protocorm containing (B) pelotons. Scale bars (A) 200 µm, (B) 100 µm.

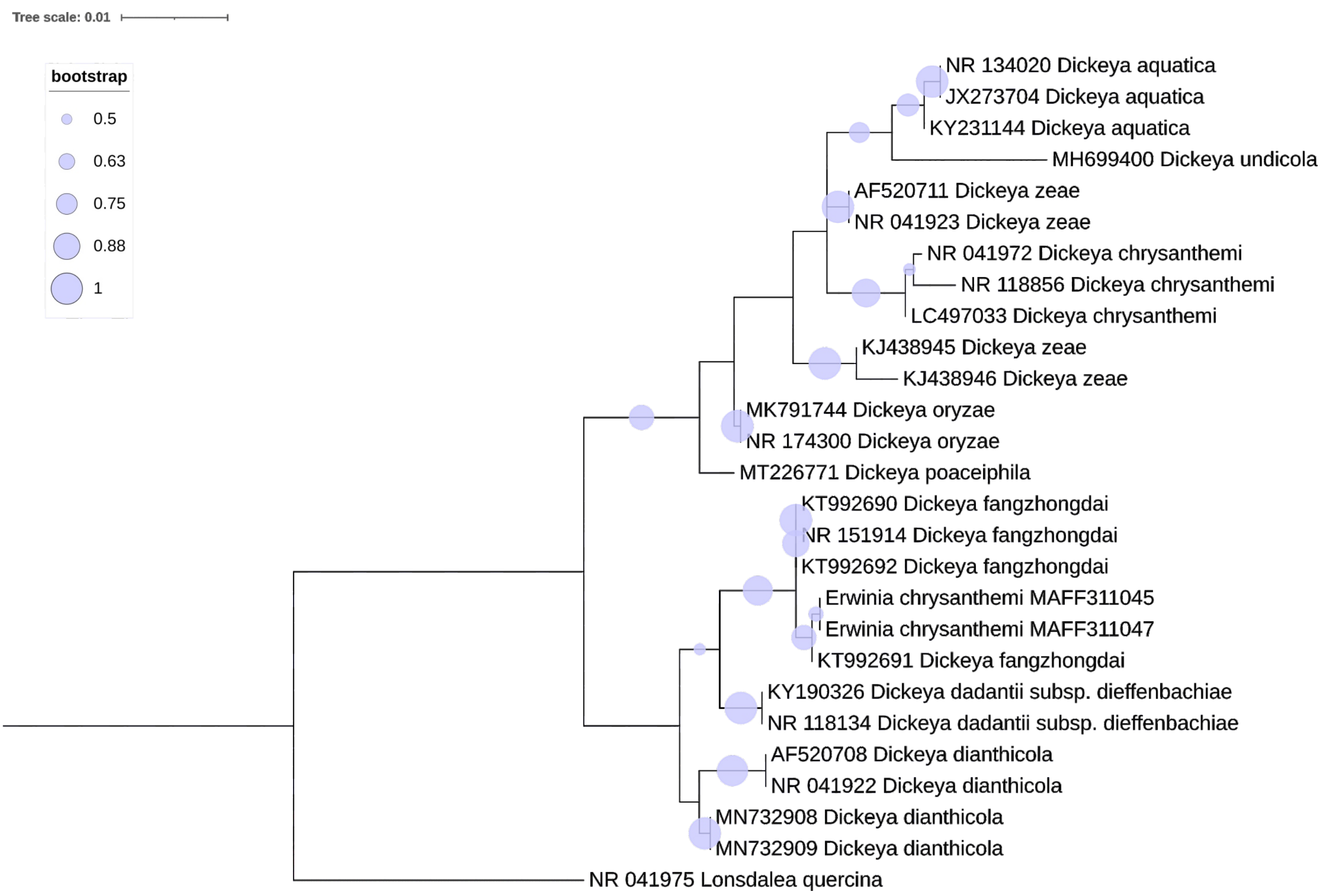


**Fig. S3** Maximum likelihood phylogenetic tree based on 16S sequences of various species within Pectobacteriaceae, including *Erwinia chrysanthemi* MAFF311045. The tree was constructed using the Hasegawa–Kishino–Yano model with gamma with invariant distribution (HKY+G+I) and 1000 bootstrap replicates. *Lonsdalea quercina* was used as the outgroup. The analysis was conducted in MEGA10 and visualized in iTOL v.6 (https://itol.embl.de).

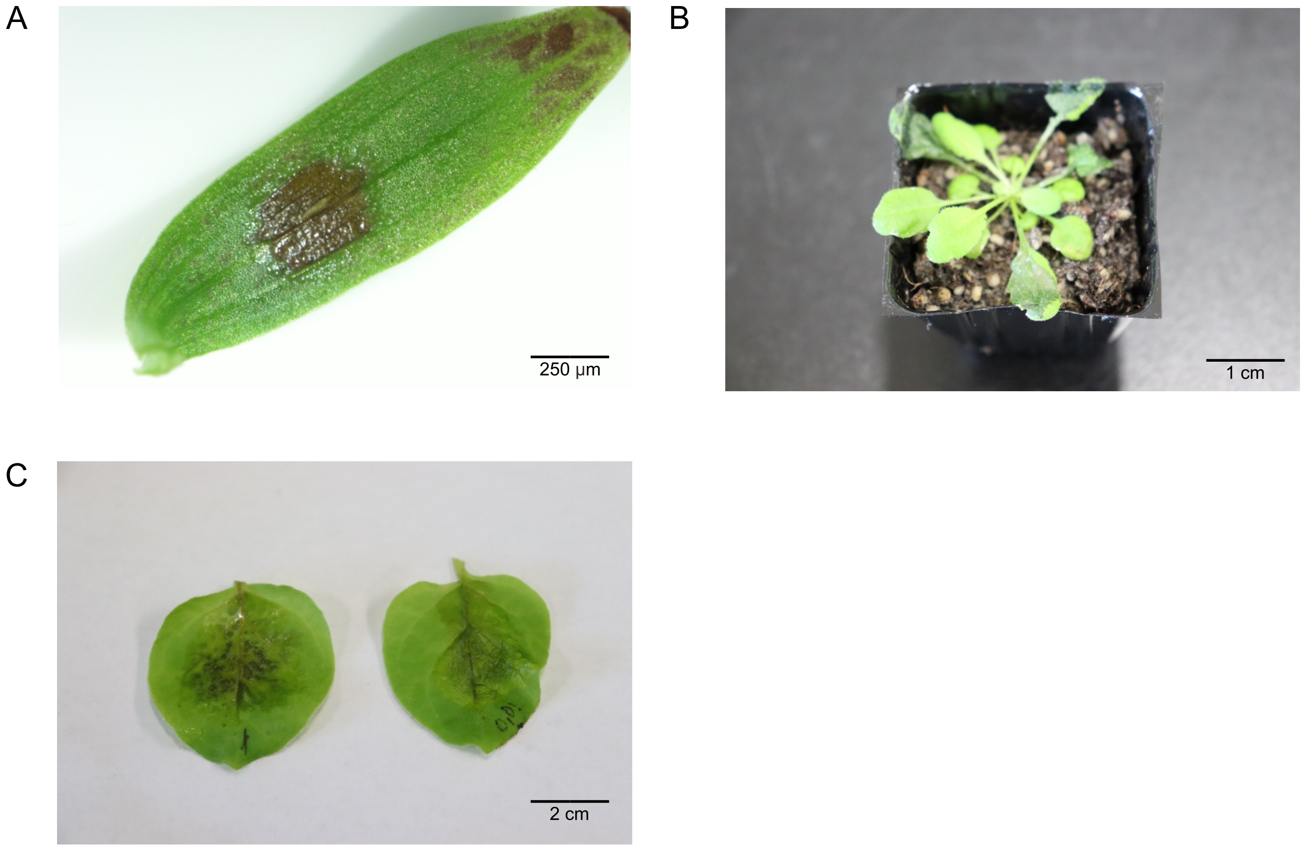


**Fig. S4** Soft rot symptoms observed in leaves of (A) *Bletilla striata*, (B) *Arabidopsis thaliana*, and (C) *Nicotiana benthamiana*.

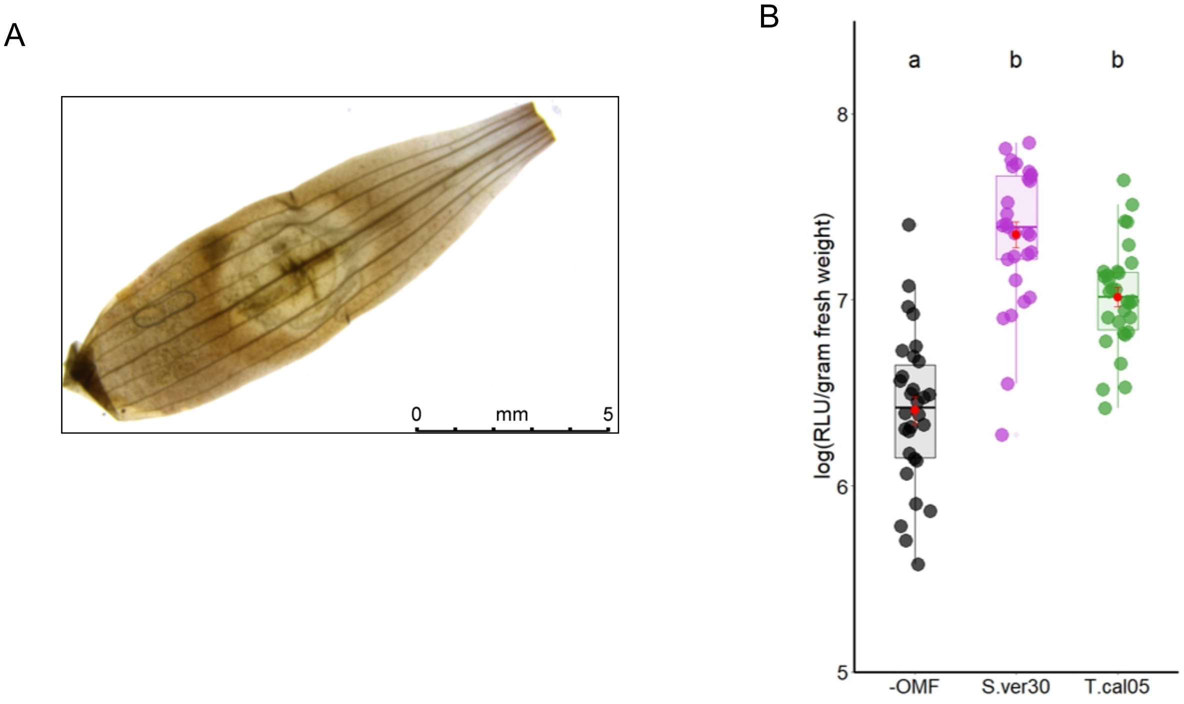


**Fig. S5** Peroxide production in infected leaves. (A) Peroxide accumulation visualized by diaminobenzidine staining (brown area). Scale bar, 5 mm. (B) Peroxide content ($p < 0.05$, Kruskal–Wallis test).

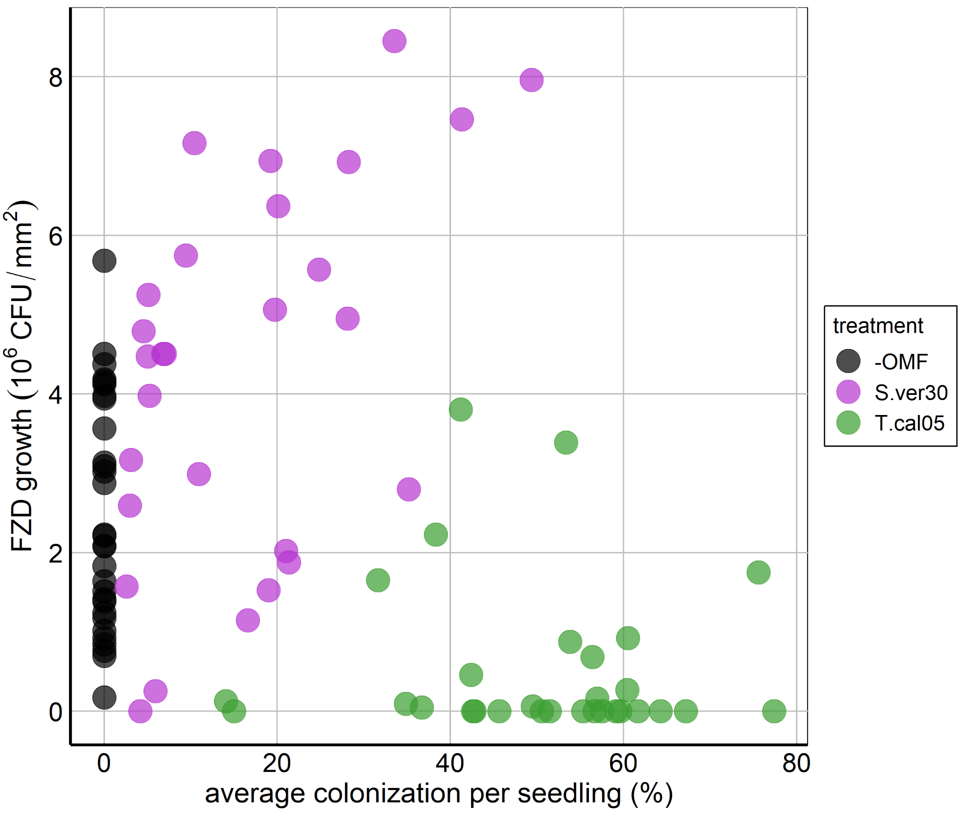


**Fig. S6** Scatter plot showing the two-dimensional relationship between OMF colonization rate and *Dickeya fangzhongdai* CFU count.

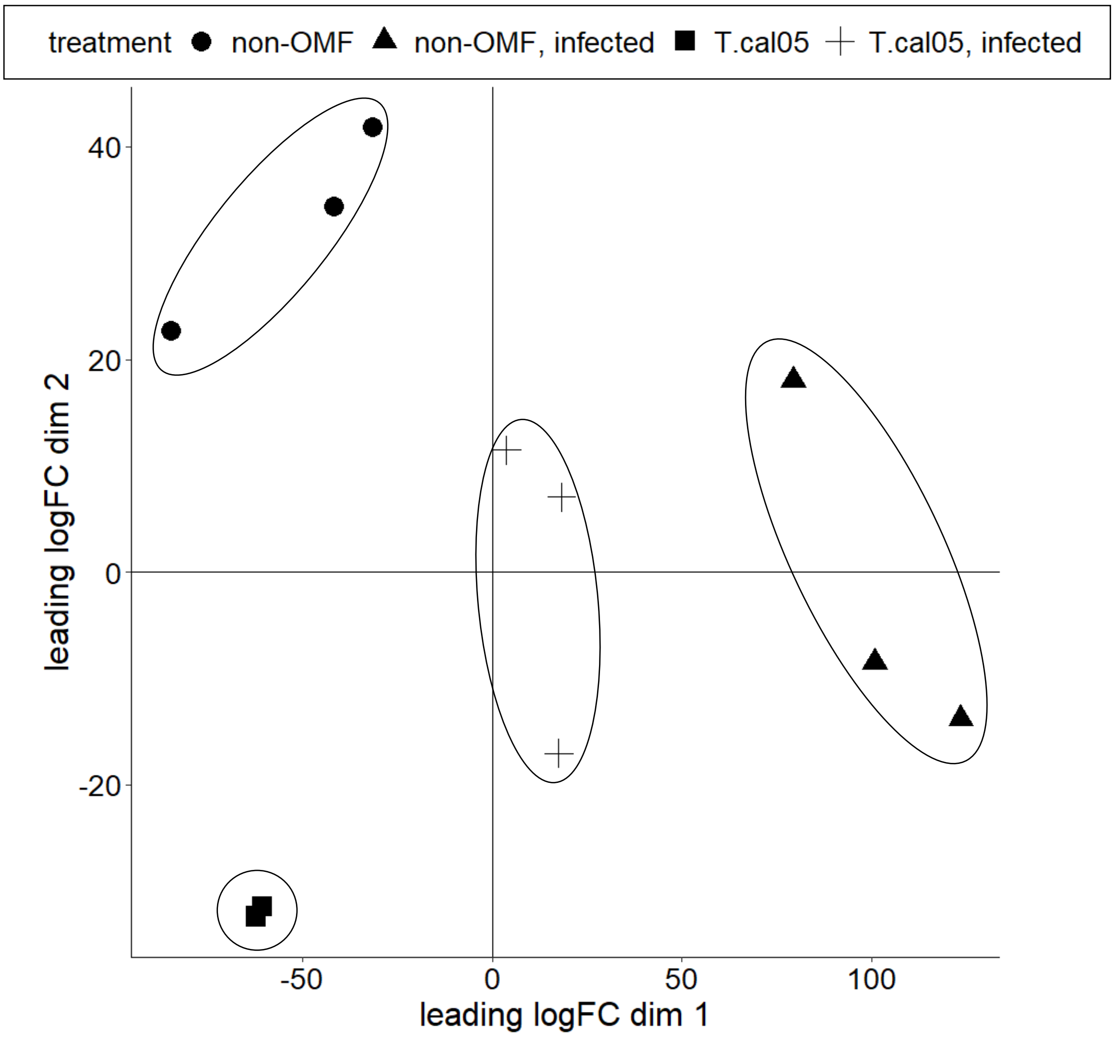


**Fig. S7** MDS plot showing separation of DEGs among all treatments.
